# Understanding boundary effects and confocal optics enables quantitative FRAP analysis in the confined geometries of animal, plant, and fungal cells

**DOI:** 10.1101/059220

**Authors:** James L. Kingsley, Jeffrey P. Bibeau, S. Iman Mousavi, Cem Unsal, Zhilu Chen, Xinming Huang, Luis Vidali, Erkan Tüzel

## Abstract

Fluorescence Recovery After Photobleaching (FRAP) is an important tool used by cell biologists to study the diffusion and binding kinetics of vesicles, proteins, and other molecules in the cytoplasm, nucleus or cell membrane. While many FRAP models have been developed over the past decades, the influence of the complex boundaries of three-dimensional cellular geometries on the recovery curves, in conjunction with ROI and optical effects (imaging, photobleaching, photoswitching, and scanning), has not been well studied. Here, we developed a three-dimensional computational model of the FRAP process that incorporates particle diffusion, cell boundary effects, and the optical properties of the scanning confocal microscope, and validated this model using the tip-growing cells of *Physcomitrella patens*. We then show how these cell boundary and optical effects confound the interpretation of FRAP recovery curves, including the number of dynamic states of a given fluorescent protein, in a wide range of cellular geometries-both in two and three dimensions-namely nuclei, filopodia, and lamellipodia of mammalian cells, and in cell types such as the budding yeast, *S. pombe*, and tip-growing plant cells. We explored the performance of existing analytical and algorithmic FRAP models in these various cellular geometries, and determined that the VCell VirtualFRAP tool provides the best accuracy to measure diffusion coefficients. Our computational model is not limited only to these cells types, but can easily be extended to other cellular geometries via the graphical Java-based application we also provide. This particle-based simulation-called the Digital Confocal Microscopy Suite, DCMS-can also perform fluorescence dynamics assays, such as Number and Brightness (N&B), Fluorescence Correlation Spectroscopy (FCS), Raster Image Correlation Spectroscopy (RICS), and could help shape the way these techniques are interpreted.

## 1. Introduction

Due to the popularity of confocal laser scanning microscopes, FRAP has emerged as a prominent technique to measure protein mobility, and throughout the past decade has appeared in over 150 publications annually [1]. During a typical FRAP experiment; a cell expressing a fluorophore of interest is subjected to a high intensity laser pulse which permanently abolishes the fluorescence properties of the fluorophore—a process called photobleaching. This laser pulse is specifically localized to a predetermined Region Of Interest (ROI) inside the cell, and over time fluorophores not subjected to the bleach move into the ROI, leading to a recovery of the local fluorescence.

The rate and directionality of this fluorescence recovery can then be used—via a model—to determine an array of important characteristics of the molecule of interest, diffusion coefficients, binding kinetics, and the number of dynamic states of the fluorophore. These properties are important for a wide array of biological questions. What is the contribution of diffusion in the transport of proteins? Is a protein part of a complex that changes its mobility? Is the protein associated with the cytoskeleton or other structures that could change its dynamics states? What are the *in vivo* kinetics of specific protein-protein interactions?

To obtain a physical constant—such as the diffusion coefficient—that can help answer these questions using FRAP, a model must be used. The two most commonly cited FRAP models are by Axelrod et al. [2] and Soumpasis et al. [3], and they use an analytically calculated recovery profile of a two-dimensional ROI inside a cell with an infinite boundary. Although these models are frequently used—over 650 combined citations to date—more recent studies have included and explored additional relevant FRAP parameters such as the finite confocal scan rate during photobleaching [4–9], arbitrary photobleaching profiles [10], confocal imaging [5], and cell shape [11–14]. Despite this wealth of analytical models, as well as algorithmic approaches [10, 13, 15–19], it remains unclear whether these models can yield accurate estimates of the diffusion coefficients for instance, when applied using realistic optical settings in actual cellular geometries. A simple thought experiment (described in detail in **Section S8 in the Supporting Material**, also **Fig. S19**) based on the method of images can show that geometric effects can produce a factor of four difference in apparent diffusion coefficient under ideal conditions. Determining these quantities accurately is important as this improves the significance of the biological conclusions derived from modeling of the cellular processes that are studied.

To provide more accuracy in analyzing FRAP recovery, interactive solutions such as the VirtualFRAP tool (part of VCell environment) [13] and simFRAP [10] have been developed, allowing the calculation of diffusion coefficients in arbitrary two-dimensional geometries. While these algorithmic approaches make fewer assumptions than existing analytical models [2, 3], they still rely on the time-scale-invariance of the process, which is not necessarily true in the case where the bleaching and imaging durations are not instantaneous. Without a comprehensive model that incorporates all the relevant aspects of FRAP, it remains a challenge to determine if a particular analytical or algorithmic method is appropriate to measure diffusion coefficients or identify dynamic states for a specific case.

While it has been shown in *silico* that cellular geometry can influence FRAP[12], this effect is often not taken into account when analyzing recovery curves obtained from arbitrary three-dimensional cellular geometries. When analytical models are unable to quantitatively describe these geometric effects, oftentimes multi-exponential fitting is used to make qualitative conclusions about the inherent dynamics. This introduction of spurious dynamic states—potentially interpreted as multiple diffusing species, active transport systems, or binding/diffusion reactions—as a result of the fitting process can further confound the problem, leading to misinterpretations of the underlying biology. These conclusions can further be reinforced by an excellent goodness of fit, regardless of the underlying physical meaning.

Given the widespread use of FRAP, there is a need for a rigorous approach—incorporating optical, geometric, and diffusive effects—that would allow accurate model selection for any cell type. Here, we developed an experimentally validated particle-based diffusion model that allows us to conduct not only FRAP, but also other fluorescence fluctuation based analyses on a wide variety of three-dimensional cellular geometries. We then quantitatively demonstrate that cell shape is one of the predominant factors that can influence FRAP recoveries, and affect measured diffusion coefficients for commonly used FRAP models. Since there is a rich parameter space that influences fluorescent recovery, it would be misleading to make absolute conclusions about FRAP recoveries in a given system. To help remedy this challenge, and to make predictions for specific experimental setups and systems, we provide a free, user-friendly, cross-platform, and GPU optimized version of our software, namely Digital Confocal Microscopy Suite (DCMS) to the readers [20].

**Figure 1.**
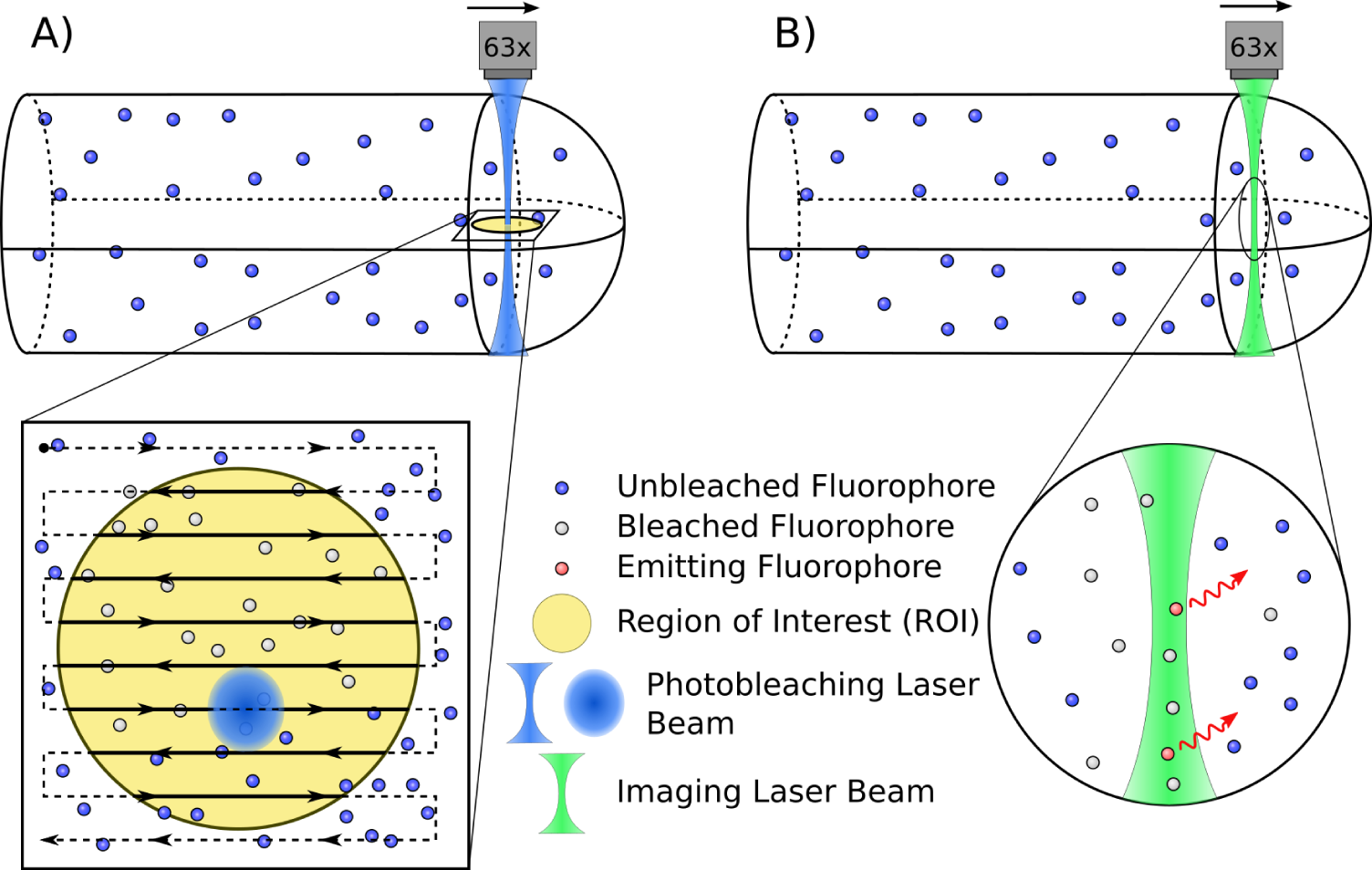
Illustration of 3D scanning confocal FRAP process. A) Scanning photobleach interacting with confined particles. Arrows indicate the bleaching pattern of the Gaussian photobleaching laser beam within the yellow circular region. B) Particle excitation and emission. Here the green diffraction limited laser scans across the image and locally excites fluorophores that emit red light. Images are not to scale.

## 2. Materials and Methods

### 2.1 FRAP Model in Digital Confocal Microscopy Suite (DCMS)

In order to accurately measure the diffusive dynamics of a given molecule in complex cellular geometries, we created a, particle-based simulation that consists of non-interacting Brownian particles, with a constant diffusion coefficient [21]. Particles are contained within a region defined by a boundary surface with reflective boundary conditions (see **Section S1.2 in the Supporting Material**). This surface can be any triangulated mesh representing the cell boundary of interest, however, when possible, we used an analytical description of the cell shape (e.g. hemisphere capped cylinder for moss) to reduce computational cost. In all of the production runs used for analysis here, we used 10^6^ particles within this simulation volume.

The simulation also incorporates properties of the optical system, i.e. imaging and bleaching (as illustrated in **Fig. 1**), finite scan rates of the confocal microscope, the PSF, and ROI size and shape related effects, as described in detail in **Section S1.3** in the **Supporting Material**. Briefly, imaging is performed by scanning across the region corresponding to the output image, using a squared Gaussian beam Point Spread Function (PSF). The experimental PSF is measured (see **Section S1.3**) and used to determine the parameters for the squared Gaussian. Photobleaching is performed by scanning across the ROI, and stochastically photobleaching fluorophores within the beam (See **Fig. S5** and **Movies S1-S2**).

Furthermore, the DCMS simulations can also conduct reversible photoswitching during imaging acquisition and bleaching events. If this reversible photoswitching is neglected, especially for 3xmEGFP, it can lead to overestimates in measured diffusion coefficients [22]. However, since the kinetics of this process can dependent on laser power [22] (see **Fig S9B**), it would add additional model parameters. In order to increase the accuracy of our results, and avoid a large parameter scan, we experimentally measured and performed the necessary corrections for acquisition photobleaching (see **Section S2.2** in the **Supporting Material**) and reversible photoswitching (see **Section S2.3** in the **Supporting Material**).

Additionally, although it was not utilized in this manuscript, DCMS also supports a number of other choices, such as alternative PSF forms, reversible photobleaching and photoswitching, and binding kinetics.

### 2.2 Cell Culture and Sample Preparation

FRAP experiments were conducted on the caulonemal cells of the moss *Physcomitrella patens*; moss tissue was cultured on cellophane placed on top of the solidified agar. Microscope samples were prepared in QR-43C chambers (Warner Instruments) as follows. First, 25 mm bottom coverslips were plasma treated for three minutes to yield a hydrophilic surface. A solution of 0.8% type VII agarose in P_p_NO_3_ medium (refer to Ref. [23] for details) was melted then added directly to the coverslips. A small cellophane square (1 *cm*^2^) with moss tissue, grown for seven days after sub-culturing, was inverted and placed onto the agarose. To obtain flat cultures, a second untreated coverslip was placed on top of the cellophane and flattened using a blunted syringe. Agarose was solidified by placing the cultures onto a surface at 12—C. Once the agarose solidified, the top coverslip was gently removed from the top of the preparation. The remaining preparation was submerged in P_p_NO_3_ and the cellophane was removed. With the moss firmly adhered to the agarose, the entire coverslip was added to the QR chamber. The chambers were capped with 18 mm coverslips and connected to silicone tubing with inner and outer diameters of 0.03 *in* and 0.065 *in*, respectively. The tubing was connected to a peristaltic pump and liquid P_p_NO_3_ was perfused through the chambers over night. Liquid P_p_NO_3_ was made two days prior to perfusion and was filter sterilized immediately before its use. During latrunculin B treatment, a solution of 10 *μM* latrunculin B in P_p_NO_3_ was perfused through the chamber for 20 minutes.

### 2.3 Confocal Imaging

FRAP experiments were conducted using a Leica TCS SP5 scanning confocal microscope and the Leica FRAP Wizard. To conduct FRAP experiments, a 63X objective was used with a numerical aperture (NA) of 1.4. In the software settings, the pinhole was set to 2.00 airy disks and the camera zoom was set to 9. Images of 256x256 pixels were acquired with a depth of 12bits. To visualize 3xmEGFP, the Argon laser was set to 75% power with a bidirectional scanning speed of 2800 Hz and the 488 nm laser line was set to 10% power in the FRAP wizard. The emission bandwidth for 3xmEGFP was set between 499 nm and 546 nm. During bleaching events, all laser lines were set to 100%.

### 2.4 FRAP Post Acquisition Processing and Analysis

Several confounding factors that can influence the interpretation of FRAP results were examined, namely, image acquisition induced photobleaching/reversible photobelaching (**Section S2.2**), reversible photo-bleaching following intentional photobleaching (**Section S2.3**), confocal detector linearity (**Section S4**), and the effects of imaging and photobleaching away from the medial cell plane (**Section S5**). We then constructed the FRAP processing scheme depicted in **Fig. S6**. This flowchart illustrates how the experimental and simulated FRAP images were processed and analyzed to yield the diffusion coefficients and highlight experimental boundary effects. Briefly, images from FRAP experiments, saved as tiff stacks, were converted into fluorescence recovery curves by averaging the intensity of the pixels within the ROI. Replicate experiments were then averaged and divided by acquisition bleaching controls. After processing the recovery curves, we used an argument minimization scheme to find the best fit simulation parameters (**Section S2.4**). Confidence intervals for the best fit parameters were found via Monte Carlo simulations (**Section S2.5**). To analyze the directionality of fluorescence recovery spatially the photobleaching ROI was cropped, corrected for the limited available volume at the cell tip (**Section S3.1**), and subjected to Fourier analysis (**Section S3.2**). A flow chart depicting this analysis can be found in **Fig. S12**.

### 2.5 Experimental Measurement of the Point Spread Function (PSF)

In order to experimentally measure the PSF of the confocal microscope, beads were used from the Invitrogen PS-Speck Microscope Point Source Kit P7220 (Thermo Fisher Scientific, Waltham, MA). Preparations were made by adding 5 *μL* of 0.01% polylysine and 5 *μL* of bead solution to a dry microscope slide. A coverslip was then placed directly onto the slide and sealed with wax. Beads were visualized and measured with the Leica TCS SP5 scanning confocal microscope, using the settings described in **Section 2.3**. Green fluorescent beads were used to match the 3xmEGFP fluorophore, and z-stacks of the 175 *nm* beads were taken. From these z-stacks, a three-dimensional reconstruction of the bead’s intensity profile was created as shown in **Fig. S5a**. These profiles were then used to determine the three-dimensional functional form and parameters of the PSF, following the procedure defined in **Section S1.3**.

### 2.6 GPU Accelerated Computation

Due to the high computational cost associated with the motion of many particles, we wrote a parallel simulation code to take advantage of General Purpose Graphics Processing Unit (GPGPU) computing. In order to provide a general purpose tool, available to a wider audience, we made a graphical-user-interface-based simulation tool (DCMS). This interface was written in Java, using the cross-platform AMD APARAPI library [24] to use OpenCL computation whenever possible. This also allows the software to efficiently use GPGPU computing when available, and fall back to CPU computing if it is not, in a cross-platform manner. Our standard production runs, using the moss geometry and 161-image, 2-bleach sequence, take approximately half an hour to run (on an AMD RX 480 GPU). More modest simulations, using fewer particles, fewer imaging steps, or more simple (fewer polygon) geometries, for example, can be performed in a few minutes.

In order to more effectively perform the large number of runs required for our library fitting approach used in **Section S2.4**, we also wrote a version of the software using NVidia’s CUDA platform. Combined with some platform-specific optimizations, as well as using an analytical form of the ideal moss geometry rather than a triangulated mesh, our run-time was reduced by approximately a factor of 10 (i.e. to 140 s on an NVidia GTX 780Ti) (see Ref. [25] for computational details).

## 3. Results

### 3.1 Cell Shape Influences Fluorescence Recoveries *in vivo*

In order to study the influence of cell shape and boundaries on fluorescence recovery *in vivo*, we conducted FRAP experiments in the moss *Physcomitrella patens*, at two different ROI locations, one at the cell edge and another at the center of the cell (as depicted in **Fig. 2b**). We processed the curves to correct for acquisition and reversible photobleaching, and ensured that our experiments were in the linear range of the detector (Sections S2.2, S2.3 and **S4**). Additionally, recoveries at the cell edge exhibited a slower rate of recovery and higher fluorescence plateau when compared to the center. To ensure that these observations were not due to filamentous actin localization at the cell edge, we performed the same analysis in the presence of an actin depolymerization agent, latrunculin B at 10*μM* (**Section S6**). Latrunculin B treatment, which has been shown to completely depolymerize the actin cytoskeleton at this concentration[26], had no influence on the observed fluorescence recovery(**Fig. S18**).

To determine if a single diffusion coefficient could reproduce the different rates of recovery observed at the edge and the center, we simulated FRAP recovery curves at the edge and center for a range of diffusion coefficients using the model described in **Section 2.1**, and **Section S1** in the **Supporting Material**. In our fitting routine (**Section S2.4**), we used one diffusion coefficient to fit recovery at both the edge and the center, which yielded a value of *D* = 8.25 ± 0.358 *μm*^2^ *s*^−1^. This diffusion coefficient is within the expected range for 3xmEGFP under physiological conditions[27], and using the Einstein-Stokes relation can be used to estimate an effective viscosity of the moss cytoplasm—approximately one order of magnitude higher than that of water in this case. The simulations at the cell edge exhibited both a slower rate of recovery and higher fluorescence plateau when compared to the center, recapitulating experimental recoveries. The slower recovery at the cell edge is due to geometrical constraints provided by the apical plasma membrane, i.e., proteins cannot flow in or exchange in all directions as they can at the center of the cell. The higher plateau at the cell edge is caused by photobleaching fewer particles at the cell edge since the three-dimensional Gaussian laser beam extends outside of the cell volume at the edge, in contrast to the fully encased beam at the cell center.

Fourier analysis of the spatial fluorescence recovery further supports the claim that cell shape can influence the interpretation of FRAP curves (Sections S3.2 and S3.3). Specifically, experimental and simulated fluorescence concentration gradients recovered in the same way, as characterized by the first mode of the Fourier series representation of the spatial recovery profile (**Fig. S13a** and **Movie S3**). Our simulation results in conjunction with experiments highlight the strong influence of boundaries on recovery, and shows that erroneous conclusions about the fraction of bound molecules and fluorescent mobility at the edge and center can be made if such geometric effects are not taken into account. It is important to note that boundary effects are masked by the additional effects (such as scan speed) associated with a fast recovery, and become more apparent at lower diffusion coefficients (**Fig. 3**). In the limit of a fast scan speed, each image or photobleach happens at a single point in time, and the curve is well sampled during recovery. However, with a slower scan speed—or a fast recovery—each image spans a large amount of the recovery time, and the resultant recovery is under-sampled, distorting the apparent fluorescence recovery.

**Figure. 2.**
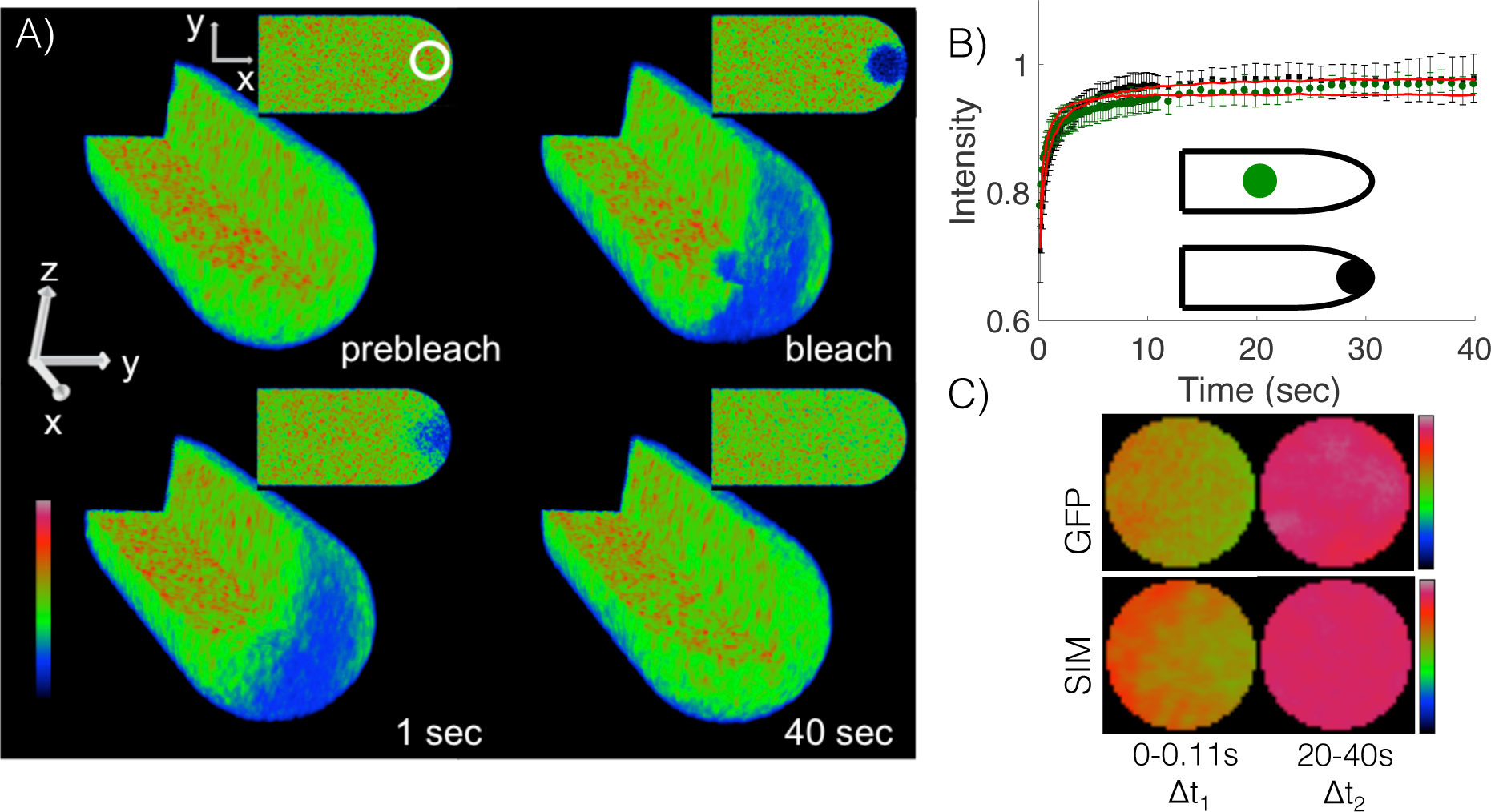
Cell shape influences fluorescence recovery *in vivo* and *in silico*. A) Three- and two-dimensional rendering of simulated scanning confocal photobleaching and recovery. Artificially fast imaging scan rates were used to illustrate three dimensional properties of the simulation. The full animated form of this panel can be found in **Movies S1 and S2**. B) Fluorescence recovery of 3xmEGFP cells at the edge (black squares) and center (green circles). Best fit simulation indicated in red. *n* = 14 and 7 for the edge and center, respectively. Error bars represent standard deviation. C) Cropped and frame averaged photobleaching ROI at the cell edge of 3xmEGFP cell line (top) and simulation (bottom). Time intervals for frame averaging are Δ*t*_1_ = 0 − 0.11 *s* and Δ*t*_2_ = 20 − 40 *s. n* = 14 and 50, respectively. ROI is 4 *μm* in diameter. Image intensity denoted with rainbow lookup table.

### 3.2 Exponential Fitting Does Not Necessarily Reflect the Underlying Dynamics

When a proper analytical model is not available to describe a given fluorescence recovery, a common practice is to fit the recovery data to a series of exponentials; the number of exponentials needed to fit the recovery has been interpreted as the number of “dynamic states” of the fluorophore [28–33]. In this context having multiple dynamic states refers to the various modes by which a fluorophore recovers during a FRAP experiment. These modes can include multiple diffusion coefficients, chemical reactions, and active transport, or any combination of the three. Specific examples would include: the presence of more than one diffusion coefficient in a reaction diffusion system; two modes of fluorescence recovery in an active transport and diffusion based system; two unique reaction rates in a reaction dominant FRAP experiment.

Fundamentally, exponential fitting should only be used to identify unique reaction rates in a reaction dominant FRAP experiment [34]. This is because diffusion-based fluorescence recovery asymptotically behaves as a power law, and neither a single, nor a double exponential can fundamentally reproduce this behavior at long times (see **Section S9**, also **Fig. S20**). Hence, there is no theoretical justification for fitting a single or double exponential to a diffusion based recovery.

As the divergence between asymptotic behavior and the exponential functional form often occurs at times longer than those that are experimentally relevant, however, one may still want to consider using it to fit early times in a recovery curve. While it may appear to provide an adequate fit in early recovery, as we will demonstrate below using a proof-by-contradiction approach, exponential fitting fails to capture the underlying dynamics and is not able to predict the number of dynamic states in a diffusion-based recovery. To start, we analyzed simulated fluorescence recoveries of a fluorophore with a single diffusion coefficient, *D* = 1 *μm*^2^ *s*^−1^ in the moss geometry. Simulations were conducted at the cell center and the cell edge. At both regions, a fit to the single exponential model exhibits periodic over- and undershooting, (**Fig. 3A**), whereas the double exponential model exhibits less periodicity for either location (**Fig. 3B**). Residual plots demonstrate periodic errors in the single exponential fit and the lack thereof in the double exponential fit (**Fig. 3C and D**). To determine if the models predict the trends of the fluorescence recovery, we performed a Wald-Wolfowitz runs test. This test determines if the residuals of the fit are random, where increased randomness suggests a better fit. The results of this test suggest (for this diffusion coefficient and given morphology) that neither the single nor the double exponential fits exhibited random residuals (null hypothesis of random residuals, p-values = 2 × 10^−8^ and 0.027, respectively). However, there is an improvement in fit quality with the double exponential function. In addition, other morphologies show an improvement for the double exponential fit that becomes significantly random **Table S4**. Sum of squared differences between the simulation and exponential fitting demonstrated that the single exponential fit exhibits an eight-fold higher sum of squared difference compared to the double exponential fit (**Fig. 3E**). Lastly, to determine if the single- or double-exponential model should be used, we performed an F-test to compare the change in sum of squares and the change in number of parameters between the two models. The results of this test predicted that the double exponential fits best with a p-value < 0.001 [35], for this morphology and diffusion coefficient.

**Table S4** further shows that similar results hold for other cell shapes and diffusion coefficients between 0.1 and 10 *μm*^2^*s*^−1^. Since the double exponential fit is comprised of two recovery phases, it is important to note how much each phase contributes to the recovery. For the curve generated at the cell center, the fast mode contributed to roughly 60% of the recovery while the slow mode contributed to about 40%, **Fig. 3F**, indicating that contributions from both modes are significant. Based on the results of this fitting procedure, one could naively conclude that the molecule simulated here has at least two dynamic states and that it must have two different modes of recovery. Such a conclusion would be incorrect, as we simulated only one diffusion coefficient.

As illustrated, at short times, exponential fitting inadequately describes the dynamic behavior of a FRAP curve, while at long times it diverges from the expected asymptotic limit. We therefore suggest that exponential forms should only be used to make comparisons of the relative rate of fluorescence recovery between curves.

### 3.3 Model Selection Should Consider the Effects of Boundaries, Dimensionality, and Initial Conditions

To explore the predictions made by existing analytical FRAP models (see **Section S7**), we simulated fluorescence recovery experiments for six different relevant cellular shapes—nuclei, filopodia and lamellipodia of mammalian cells, budding yeast, *S. pombe*, and tip-growing plant cells (moss)—for a range of biologically relevant diffusion coefficients (**Fig. 4A and b** and **Movies S4-S8**). We used four notable analytical models [2–4, 36] to fit to our simulated recoveries. Because of their substantial citation history, we selected both the analytical models from Axelrod et al. [2] and Soumpasis et al [3]. For its simplicity, and because it incorporates boundaries, we selected a one-dimensional analytical solution with reflective boundary conditions [36]. Because it takes into account the PSF and confocal scanning during photo-bleaching, we chose the analytical solution given by Braeckmans et al. [4]. Once the simulated recoveries were fit with these models, we measured the percent difference between the simulated input diffusion coefficient, *D*_true_, and the model predicted diffusion coefficient, *D*_model_, i.e, |(*D*_true_ − *D*_model_)/*D*_true_|*100.

The analytical models from Axelrod [2], Soumpasis [3], and Braeckmans [4], most accurately predicted diffusion coefficients when photobleaching was conducted at the cell center. As expected, all three models failed to predict diffusion coefficients reliably in long thin tubular geometries (with percent differences greater than 50% for most cases), i.e. *S.pombe* and filopodia, as shown in **Table I**. These models also exhibited similar failings in the remaining cell shapes when FRAP was conducted at the cell edge (with percent differences greater than 50% for most cases). Further evidence for this spatial effect was demonstrated by conducting photobleaching and model fitting at a varying distance away from the cell edge. Consistent with strong boundary effects, as the bleaching event (ROI) was moved further from the cell edge, we observed an increased rate of fluorescence recovery and improved model predicted diffusion coefficients (see **Fig 5**). This cautions against using indiscriminately, analytical models that make infinite boundary assumptions in complex cellular geometries. These boundary effects can be further illustrated using the one-dimensional strip FRAP model that incorporates boundaries. This model predicts that bleaching at the boundary should recover with an effective diffusion coefficient 4 times slower than a centered bleach (see **Section S8**).

The one-dimensional strip FRAP was found to be the most accurate (with percent differences less than 25% in most cases) in both *S.pombe* and filopodia where the geometries closely resembled a long thin tube relative to the imaging ROI, as shown in **Table I**. Furthermore, because the model accounts for boundaries, it accurately predicts diffusion coefficients at the ends of the cells. Although moss is also a long tubular cell, the strip FRAP model could not accurately measure diffusion within this cell type. This is because moss is large relative to the PSF such that the bulk of the fluorescence recovery does not happen along one dimension. The one dimensional strip model failed to measure diffusion coefficients accurately for the remaining more complicated cell shapes.

**Figure 3.**
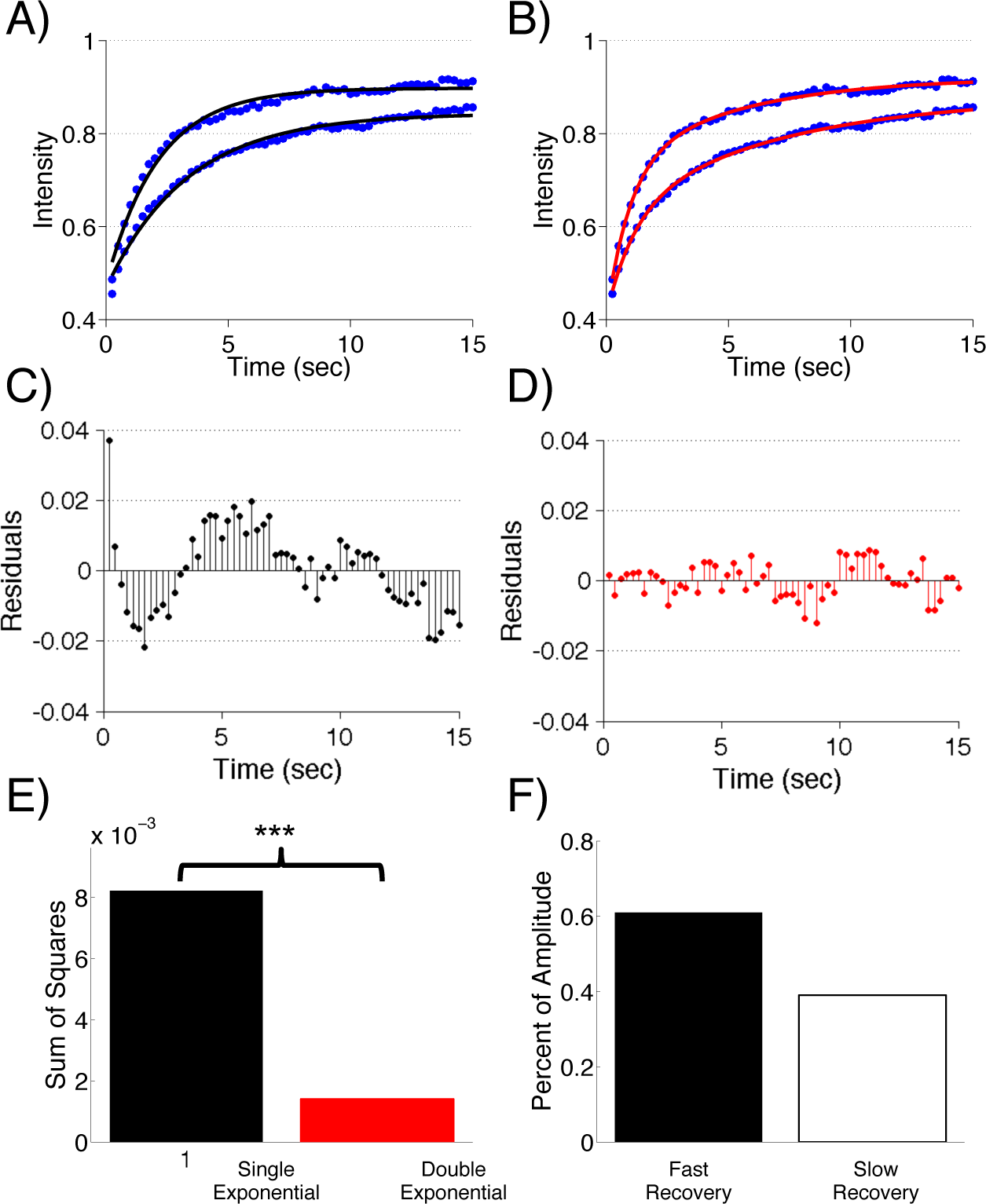
Multiple exponential fits cannot evaluate the number of dynamic states of a protein. A) Simulated moss fluorescence recovery (blue circles) with corresponding single exponential fits (black lines), D=1 *μm*^2^*s*^−1^. Photobleaching was performed at the cell edge (bottom) or in the middle of the cell (top). B) Simulated moss fluorescence recovery (blue circles) with corresponding double exponential fits (red lines), D=1 *μm*^2^*s*^−1^. Photobleaching was performed at the cell edge (bottom) or in the middle of the cell (top). C) Residuals for the single exponential fit to simulated moss recovery away from the cell boundary. D) Residuals for the double exponential fit to simulated moss recovery away from the cell boundary. E) Sum of squares for the single exponential (black) and the double exponential (red) fits. Fits were performed on the simulated moss recovery at the middle of the cell. *** indicates p-value < 0.001 of extra sum of squares F-test. F) Percent of fluorescence recovery dictated by the fast (black) and slow (white) terms in the double exponential fit. Fit was performed on moss recovery at the middle of the cell.

Lastly, we chose the VirtualFRAP tool (part of VCell environment) [13] as an algorithmic example because it incorporates the cellular boundaries, albeit in two-dimensions, and calculates a diffusion coefficient after an iteration procedure. Note that the VirtualFRAP tool is a two-dimensional continuum approach that is best suited for low NA lenses. This low NA is necessary to bleach a cylindrical region encompassing the height of the cell. The algorithmic VCell VirtualFRAP tool accurately estimated diffusion coefficients (within a 25% difference) for all morphologies with only one exception(see below), (**Table I**). This level of accuracy is consistent with our simulated bleached region being close to a cylinder as shown in **Section S11**. It also indicates that incorporating the two dimensional geometry is a good approximation for many cell types and our measured optical settings.

**Figure. 4.**
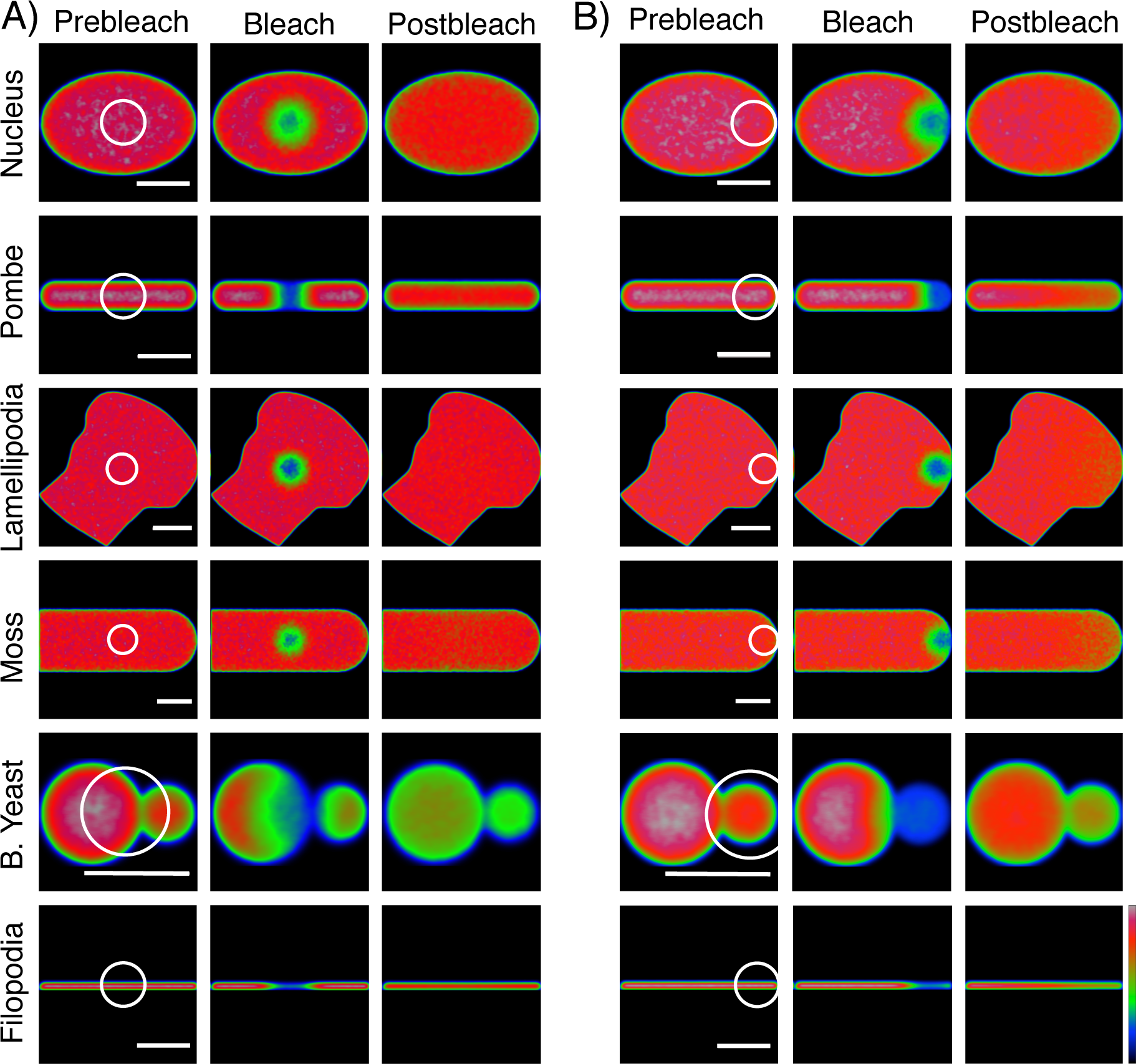
Example simulations of relevant three-dimensional cell shapes. A+B) Medial section of simulated fluorescence recoveries with photobleaching at the middle (A) or edge (B) of the cell. Post-bleach image is 16 seconds after photobleaching. Fluorescence intensity is indicated by the rainbow lookup table. Scale bar is 5*μm*.

One trend was observed across all the models, specifically that the models mostly predicted inaccurate diffusion coefficients when *D*_true_ = 10 *μm*^2^*s*^−1^. This failure is the product of the finite scan rate used during photobleaching and image acquisition, and can be attenuated at artificially fast scan speeds, **Table S5**. At 2800 Hz, a protein diffusing at 10 *μm*^2^*s*^−1^ is expected to travel roughly 80 *nm* during a single line scan. Given our 100 *nm* line spacing, this suggests that diffusion is too fast for our acquisition speed. Furthermore, this demonstrates that at the upper scan speed of our confocal system (2800 Hz), we cannot reliably measure fast diffusion coefficients greater than 10 *μm*^2^*s*^−1^, and highlights how the finite scan speed of the confocal must be considered as a limiting factor in performing FRAP experiments.

### 3.4 Membrane FRAP

FRAP analysis of molecules associated with the plasma membrane has been used extensively to characterize their dynamics [37–41]. Compared with intracellular molecules, using FRAP to estimate diffusion coefficients in membranes is simplified due to the two-dimensional nature of the membrane, as well as the slower dynamics of the molecules. To explore the influence of membrane curvature on fluorescence recovery, we simulated photobleaching experiments inside a 200 *nm* thick moss shell (**Fig. 6A**). The shell was oriented such that the imaging plane was parallel to the flat circular surface of the cylinder (**Fig 6B**). Imaging and bleaching were then conducted on the flat circular surface of the cylinder or the extreme cell apex, as shown in **Fig 6A and B**. To simulate typical membrane diffusion, *D* = 0.1 *μm*^2^*s*^−1^ was chosen. Analysis of our simulation results showed an increase in fluorescence recovery at the cell apex (**Fig 6B**).

**Figure. 5.**
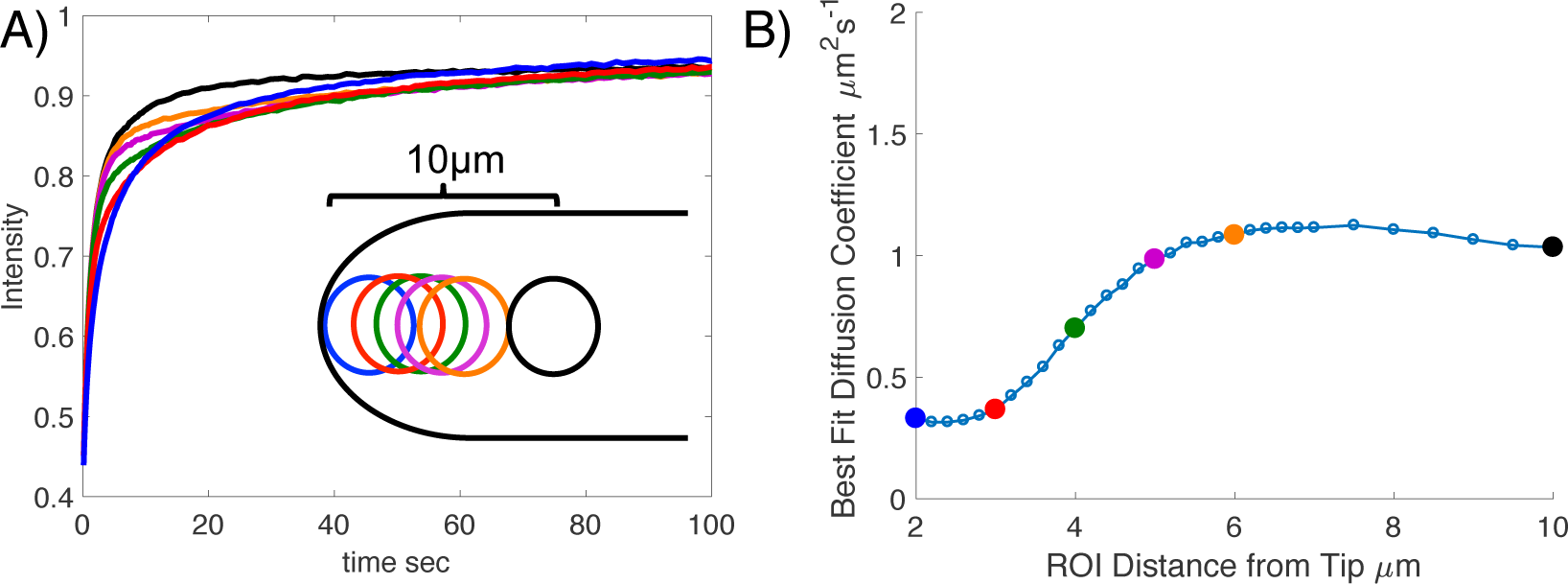
ROI Positional Dependence. A) Simulated fluorescence recovery for an array of photobleaching regions. Color of the curve denotes position of bleach as depicted. B) Best fit diffusion coefficients for the simulated recoveries in (A) when fit to the Soumpasis [3] model. Smaller dots are intermediate points, not depicted in the in (A).

**Table I.** Best fit diffusion coefficients for the four analytical models and the VCell VirtualFRAP tool. ROI is either at the cell Edge (E) or Center (C). ± represents standard error of ten simulations. Bolded text indicates cases where the model produced an answer within 25% of the true D value. Note that similar results for *D* = 0.1 *μm*^2^*s*^−1^ are given in **Table S3**. To quantify the effects of confocal scan speed, results for *D* = 10 *μm*^2^*s*^−1^, with a 280 kHz scan speed, can be found in **Table S5**. * indicates cases in which switching to a faster scan speed brought the measured value to within 25% of the true D value.

| Cellular Geometry | | Diffusion Coefficient ($\mu\text{m}^2\text{s}^{-1}$) | | | | | |
| --- | --- | --- | --- | --- | --- | --- | --- |
|  |  | Input D | Axelrod 1976b | Soumpasis 1983 | Braeckmans 2003 | 1D FRAP | VirtualFRAP |
| B Yeast | E | 1 | $0.123 \pm 0.001$ | $0.103 \pm 0.001$ | $0.103 \pm 0.001$ | $0.820 \pm 0.004$ | **$1.22 \pm 0.01$** |
| B Yeast | C | 1 | $1.95 \pm 0.02$ | $1.82 \pm 0.02$ | $1.82 \pm 0.02$ | $3.98 \pm 0.02$ | **$0.908 \pm 0.004$** |
| B Yeast | E | 10 | $3.16 \pm 0.03$ | $2.90 \pm 0.02$ | $3.25 \pm 0.16$ | **$9.44 \pm 0.04$** | **$11.7 \pm 0.1$** |
| B Yeast | C | 10 | $96.7 \pm 1.8$ | $84.5 \pm 0.3$ | $49.1 \pm 0.5$ | $44.4 \pm 0.7$ | **$8.42 \pm 0.13$** |
| Lamellipodia | E | 1 | $0.534 \pm 0.007$ | $0.530 \pm 0.007$ | $0.530 \pm 0.007$ | $6.18 \pm 0.08$ | **$1.03 \pm 0.01$** |
| Lamellipodia | C | 1 | **$0.962 \pm 0.010$** | **$0.942 \pm 0.010$** | **$0.942 \pm 0.010$** | $3.54 \pm 0.05$ | **$1.02 \pm 0.01$** |
| Lamellipodia | E | 10 | $2.16 \pm 0.04$ | $2.33 \pm 0.04$ | $2.33 \pm 0.04$ | $25.1 \pm 0.4$ | **$9.95 \pm 1.29$** |
| Lamellipodia | C | 10 | $12.8 \pm 0.6$ * | **$12.1 \pm 0.5$** | **$12.1 \pm 0.5$** | $65.6 \pm 1.3$ | **$10.0 \pm 0.2$** |
| Nucleus | E | 1 | $0.358 \pm 0.008$ | $0.351 \pm 0.008$ | $0.351 \pm 0.008$ | $3.48 \pm 0.07$ | **$1.03 \pm 0.02$** |
| Nucleus | C | 1 | $1.56 \pm 0.02$ | $1.55 \pm 0.02$ | $1.55 \pm 0.02$ | $7.18 \pm 0.14$ | **$1.01 \pm 0.01$** |
| Nucleus | E | 10 | $3.17 \pm 0.05$ | $3.38 \pm 0.05$ | $3.38 \pm 0.05$ | $23.3 \pm 0.3$ | $13.9 \pm 0.2$ * |
| Nucleus | C | 10 | $81.7 \pm 0.7$ | $46.4 \pm 0.4$ * | $40.3 \pm 0.3$ | $83.3 \pm 1.4$ | $6.36 \pm 0.13$ * |
| Pombe | E | 1 | $0.152 \pm 0.001$ | $0.135 \pm 0.002$ | $0.135 \pm 0.002$ | $1.39 \pm 0.02$ | **$1.05 \pm 0.01$** |
| Pombe | C | 1 | $0.374 \pm 0.002$ | $0.359 \pm 0.002$ | $0.359 \pm 0.002$ | **$1.17 \pm 0.01$** | **$0.948 \pm 0.004$** |
| Pombe | E | 10 | $0.994 \pm 0.005$ | $0.999 \pm 0.005$ | $2.28 \pm 0.14$ | **$11.6 \pm 0.1$** | **$11.3 \pm 0.1$** |
| Pombe | C | 10 | $6.94 \pm 0.12$ | $6.13 \pm 0.08$ | $7.40 \pm 0.12$ | $37.3 \pm 0.2$ | **$10.1 \pm 0.1$** |
| Moss | E | 1 | $0.467 \pm 0.012$ | $0.469 \pm 0.014$ | $0.469 \pm 0.014$ | $5.68 \pm 0.19$ | **$1.10 \pm 0.02$** |
| Moss | C | 1 | **$1.06 \pm 0.03$** | **$1.07 \pm 0.03$** | **$1.07 \pm 0.03$** | $4.31 \pm 0.15$ | **$1.05 \pm 0.02$** |
| Moss | E | 10 | $2.26 \pm 0.27$ | $2.43 \pm 0.29$ | $2.43 \pm 0.29$ | $32.7 \pm 3.8$ | **$12.1 \pm 0.2$** |
| Moss | C | 10 | $6.62 \pm 0.24$ | $6.98 \pm 0.23$ | $6.98 \pm 0.23$ | $52.9 \pm 2.0$ | **$7.81 \pm 0.14$** |
| Filopodia | E | 1 | $0.147 \pm 0.001$ | $0.108 \pm 0.001$ | $0.108 \pm 0.001$ | **$1.09 \pm 0.01$** | **$1.03 \pm 0.01$** |
| Filopodia | C | 1 | $0.420 \pm 0.003$ | $0.402 \pm 0.003$ | $0.402 \pm 0.003$ | **$1.12 \pm 0.01$** | **$1.03 \pm 0.01$** |
| Filopodia | E | 10 | $0.982 \pm 0.027$ | $0.978 \pm 0.030$ | $3.17 \pm 0.07$ | **$9.56 \pm 0.26$** | **$10.3 \pm 0.1$** |
| Filopodia | C | 10 | **$8.30 \pm 0.05$** | **$7.66 \pm 0.04$** | **$7.66 \pm 0.04$** | $32.0 \pm 0.2$ * | **$10.3 \pm 0.1$** |

**Figure. 6.**
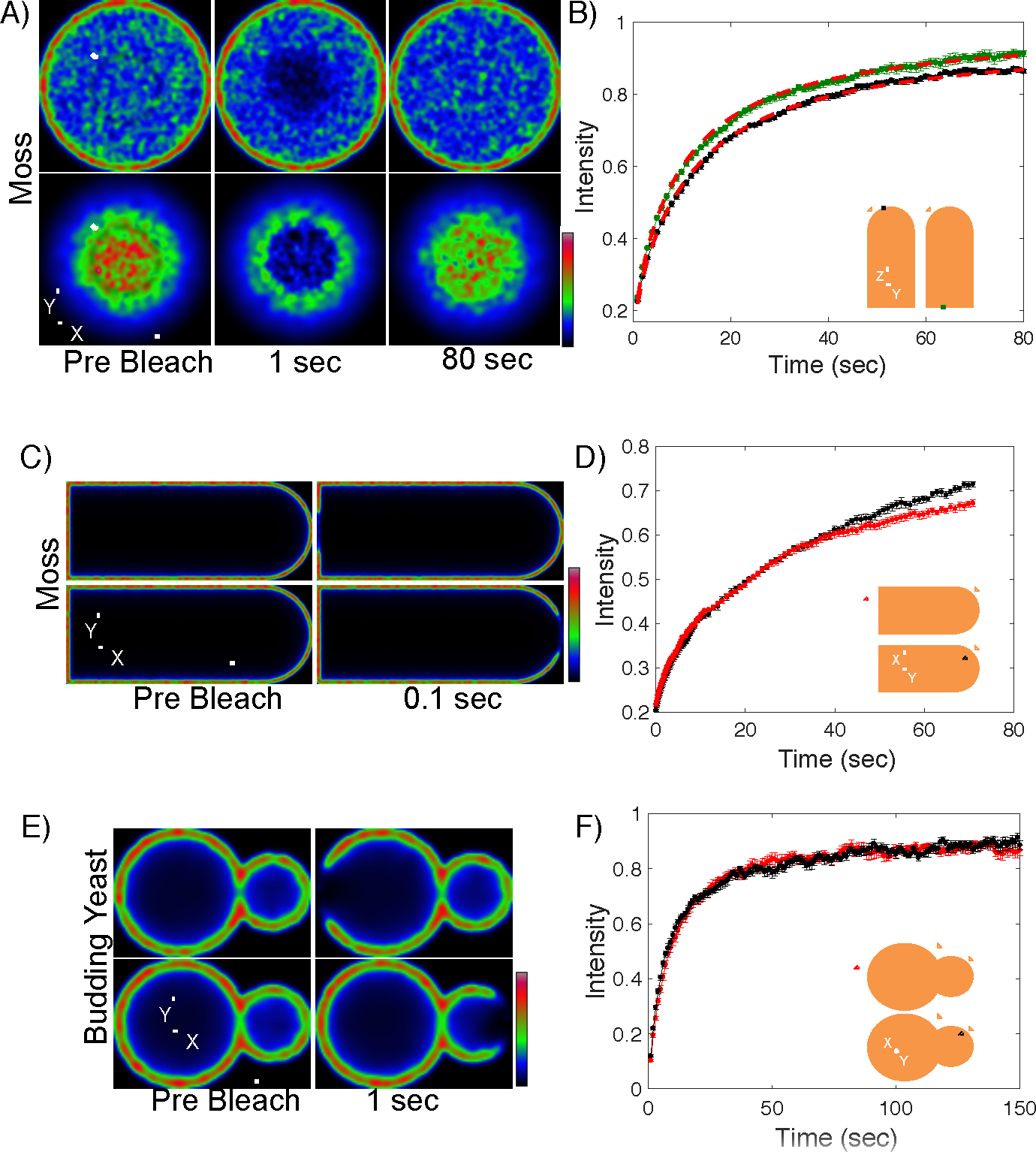
Membrane FRAP. A) Simulated fluorescence recovery images of the moss cell membrane, at the base of the cell (top) or the extreme cell apex (bottom) with bleaching and imaging planes perpendicular to the long axis of the cell. Scale bar is 3 *μm*. B) Simulated fluorescence recovery curves at the base of the cylinder (green) or the extreme cell apex (black). n=10, error bars indicate standard error. Dashed lines represent fits to the Soumpasis model (found in **Section S7**). C) Simulated fluorescence recovery images of the moss cell membrane, with bleaching and imaging planes parallel to the long axis of the cell. Scale bar is 5 *μm*. D) Simulated fluorescence recovery curves at the base of the moss cell cylinder (red) or the extreme cell apex (black). n=10, error bars indicate standard error. E) Simulated fluorescence recovery images of the budding yeast membrane, with bleaching and imaging planes parallel to the long axis of the cell. Scale bar is 3 *μm*. F) Simulated fluorescence recovery curves at mother cell (red) and the daughter cell (black). n=10, error bars indicate standard error.

When fit with the Soumpasis model (see **Section S7**), we found the measured diffusion coefficients to be *D* = 0.11 ± 0.01 and 0.13 ± 0.01 *μm*^2^*s*^−1^ at the base of the cylinder and the cell apex, respectively. This reduction in predicted diffusion coefficient can be attributed to performing a planar bleach scan on a curved membrane. During this planar scan, the membrane curvature exposes more bleachable surface area, creating an larger effective ROI. We expect this effect to become more dramatic in instances where both the ROI and membrane curvature are large.

To investigate the effects of FRAP on membranes in other cellular orientations, we simulated imaging and bleaching planes that were parallel to the long axis of the moss cell (**Fig. 6C and D**). Simulations were then subjected to photobleaching at the base of the cylinder or the extreme cell apex. At early times flourescence recoveries at both locations appeared identical, however, at long times we observed separation between the curves (**Fig. 6D**). This is beaeuse photobleaching in this orientation bleached more molecules at the base of the cylinder when compared to the apex.

To explore FRAP, in a cellular geometry often used to study membrane dynamics, we simulated photobleaching on the membrane of budding yeast (**Fig. 6E and F**). We found that photobleaching at the daughter cell or the mother had little to no influence on flourescence recoveries. Once cell orientation and effective ROI sizes are accounted for, membrane curvature has minimal influence on fluorescence recoveries, consistent with earlier modeling work[41].

## 4. Discussion

In this paper, we experimentally showed, to the best of our knowledge for the first time, the influence of cell boundaries in FRAP recovery. Our results were recapitulated using a comprehensive FRAP model that not only takes into account three-dimensional cellular geometry, but also confocal microscope optical properties. We observed that when performed close to boundaries, for instance when studying tip growing cells, fluorescence recovery is significantly slowed even though a molecule’s diffusion coefficient remains the same. As we have also demonstrated with an analytical model in **Section S8**, this results in a smaller effective diffusion coefficient. Therefore, whenever possible, we recommend avoiding FRAP closer than one ROI distance from the edge. In addition, when FRAP is conducted close to the cell boundary, the three-dimensional Gaussian laser beam may extend into the area outside of the cell volume and, as a consequence, bleach fewer fluorophores than photo-bleaching experiments performed at the cell center. These results suggest that comparison of bound fractions of molecules near cellular boundaries should be performed with caution.

The FRAP models we used from Axelrod *et al*., Soumpasis *et al*. and Braeckmans *et al*. give very similar results and are useful to obtain diffusion coefficients in flat structures such as lamellipodia and membranes. These models can also be useful for larger structures, such as plant tip growing cells or other cylindrical structures, as long as appropriately fast image acquisition rates are used relative to the diffusion coefficient in question. Our results show that for thin and elongated structures, similar to a filopodium, pombe, or an elongated fungal hypha, a simple 1D FRAP model could be sufficient, as long as it incorporates boundaries. For more complex morphologies, sophisticated approaches are necessary, such as the VCell VirtualFRAP tool. We found that the VirtualFRAP tool—an algorithmic approach—performs well in all of our simulated cases, when an appropriately fast scan rate is used, with less than 25% error in estimation of the diffusion coefficient. Importantly, VirtualFRAP performs well at boundaries, but when two-dimensionality cannot be approximated (i.e. when high NA is required), it might becomes necessary to apply a full optical and geometrical simulation method such as the one presented in here. This level of accuracy, however, is often times adequate, and will also allow for the estimation of the effective viscosity, and can provide further insight into the cytoplasmic forces.

Overall, we found that the analytical models considered in this paper can generally fail if the underlying assumptions do not match the cellular geometry or confocal optical properties that were used. While this is expected, given the misuse of these models in the literature, we recommend that they should not be used naively without consideration of the experimental setup. This caveat also applies to our DCMS software, as it makes assumptions about the properties of the system (simple diffusion, specific PSF shape, finite fluorophore reservoir, etc.), as well as requiring that the researcher knows and inputs the parameters of their experimental setup.

In addition, we find that extraneous recovery states resulting from fitting a series of exponentials can be explained by the geometrical and optical properties of the system, rather than an underlying biological mechanism. Moreover, when using analytical FRAP diffusion/binding kinetics models, we expect these effects to influence binding and dissociation constants. We additionally discourage the use of exponential fitting without theoretical justification, as its asymptotic behavior does not match that of a primarily diffusive system. A detailed analysis of these effects is beyond the scope of this work, and is left for future studies.

Although our analysis showed how boundary and optical effects can confound FRAP analysis, these conclusions cannot be generalized to every experimental setup, and further validation across additional models is required. In order to make this model validation process more accessible, for experimentalists using existing models or theorists developing new analytical models, we developed an interactive graphical interface for the Java-based version of our simulation, called Digital Confocal Microscopy Suite (DCMS). This tool allows the users to simulate specific experimental conditions—cell type, microscope settings, and diffusion coefficient—and easily generate simulated recovery curves. Any given model (analytical or algorithmic) can then be fit to these simulated recovery profiles to calculate the model predicted diffusion coefficient. The difference between the model predicted and input (to the simulation) diffusion coefficients can then be used to determine the validity of the model of interest. DCMS can also serve as a useful and freely available [20] teaching tool, not only to explore optical and boundary effects in FRAP, but also to demonstrate how scanning confocal microscopes can be used in fluorescence dynamics techniques, such as Raster Image Correlation Spectroscopy (RICS) [42], Number and Brightness (N&B) [43, 44], and Fluorescence Correlation Spectroscopy (FCS) [45].

## Author Contributions

JLK developed the simulations and JPB performed the experiments. JLK and JPB analyzed the data, wrote the manuscript. ZC, XH, JLK contributed to the initial CUDA implementation of the simulations. CU, SIM, and JLK developed DCMS APARAPI implementation, SIM wrote the corresponding Java Graphical User Interface (GUI) for DCMS. ET and LV conceived the study, wrote the manuscript and supervised the overall study.

## Acknowledgments

We thank Fabienne Furt for development of some of the cell lines used. We also thank Leica Microsystems GmbH for technical guidance, and all members of the Tuzel and Vidali Labs for helpful discussions. We also would like to thank Dr. William Hancock (Penn State University) for careful reading of the manuscript and his insightful suggestions. This work was supported by National Science Foundation CBET 1309933, and NSF-MCB 1253444. ET and LV also acknowledge support from Worcester Polytechnic Institute Startup Funds. JLK acknowledges support from the WPI Alden Fellowship. We thank anonymous reviewers for their constructive comments.

## Supporting Citations

References [46–57] appear in the **Supporting Material**.

## SUPPLEMENTARY MATERIAL

### S1. FRAP Model in Digital Confocal Microscopy Suite (DCMS)

#### S1.1 Diffusion of Particles

We used Brownian dynamics to simulate the diffusion of fluorescent particles in three-dimensional cellular geometries [1, 2]. The particles are assumed to be in a fluid with a very low Reynolds number, in which inertial effects are negligible, and the Stokes equation, i.e.,

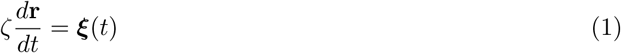

applies for each particle. Here *ξ_i_*(*t*) is the *i*^th^ sample of random Gaussian noise with the properties

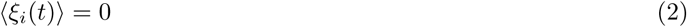

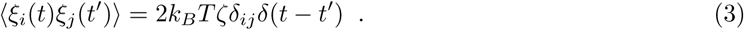

Assuming spherical particles with radius *R*, the friction coefficient *ζ*, is given by

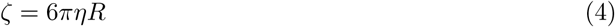

for the dynamic viscosity *η*. We can relate this *ζ* to the diffusion coefficient *D*, using the Einstein relation

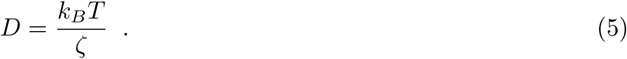

#### S1.2 Boundary Conditions

The particles in the simulation were constrained to stay within a region defined by a triangulated mesh, as shown in **Fig. S1a**. The constraint was imposed by adding an elastic collision between the particles and the boundary of the geometry, displayed in **Fig. S1b** and **S1c**. For each particle, where *d_i_* is the attempted displacement before collision, during each time step, a line segment is drawn from the previous position along its direction of travel. For each of the triangles that compose the boundary, the line segment connecting the current to the next position is tested to check if it intersects the triangle. Out of the intersections that are found, the first one reached is chosen as the collision to perform. The position of the particle is set to this intersection point, and the amount of extra distance is recorded. The displacement vector is then reflected along the surface normal 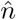, and the final displacement **d***_f_* is determined from the initial displacement **d**_*i*_ by

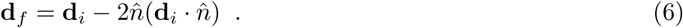

The particle then repeats the movement process (including additional collisions, if appropriate) using the remaining amount of distance.

We discretize this model using a basic Eulerian integration scheme with a variable time step. As we only need particle coordinates when we use them for imaging or photobleaching, we can choose the time step size (*δt*) to be variable. At every time step, the difference between the current time, and the time of the next imaging or photobleaching event, is calculated. This time difference is then used as the time step size for that step. It has been shown that [3] error associated with Brownian dynamics simulations with reflective boundary conditions is on the order of 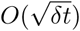. In order to keep a bound on our simulation discretization error, we place a maximum value on the possible duration of a time step. If a given time step would be larger than that maximum duration, time steps of that maximum duration are taken instead, until the time of the next event is reached. This maximum is chosen to limit the average distance that a particle can travel in a single time step, via the relation *Dδt* < 0.1 *μm*^2^. To show that our simulations are performed in the region where that error is small, we varied the maximum allowable time step size, and found that the resulting FRAP recovery curve is unaffected, as shown in **Figure S2**.

**Figure S1.**
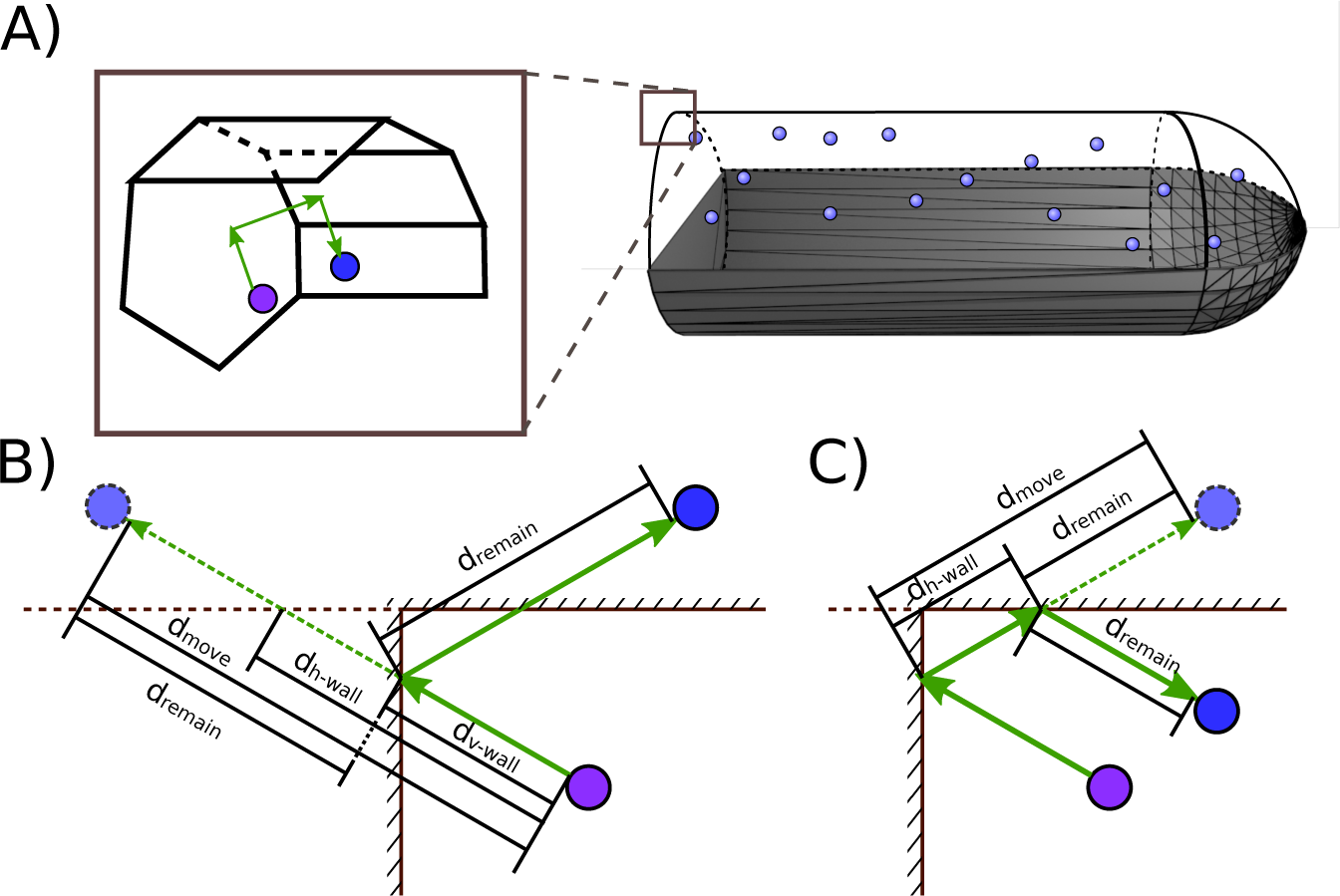
Fluid model boundary conditions. A) Visual representation of the model moss geometry. B) Example of the wall collision process for a particle attempting to move outside a square corner in two dimensions. The particle’s tentative final position after moving *d*_move_ is outside both the horizontal and vertical boundaries. This causes both distances *d*_v-wall_ and *d*_h-wall_ to be calculated. In this case, *d*_v-wall_ is the smaller, and thus happened first, so it is the chosen collision. The distance *d*_remain_ = *d*_move_ − *d*_v-wall_ is calculated, the displacement vector is reflected by the boundary, and a new tentative position is proposed a distance *d*_remain_ along that vector. C) Again the same particle’s tentative position is tested, and found to be outside the horizontal boundary. There is only one distance, *d*_h-wall_, so *d*_remain_ = *d*_move_ − *d*_h-wall_ is the only option. The displacement vector reflection process is performed again, and the particle moves its remaining *d*_remain_, arriving at yet another tentative position. As this position is inside all boundaries, it is accepted as the final position.

**Figure S2.**
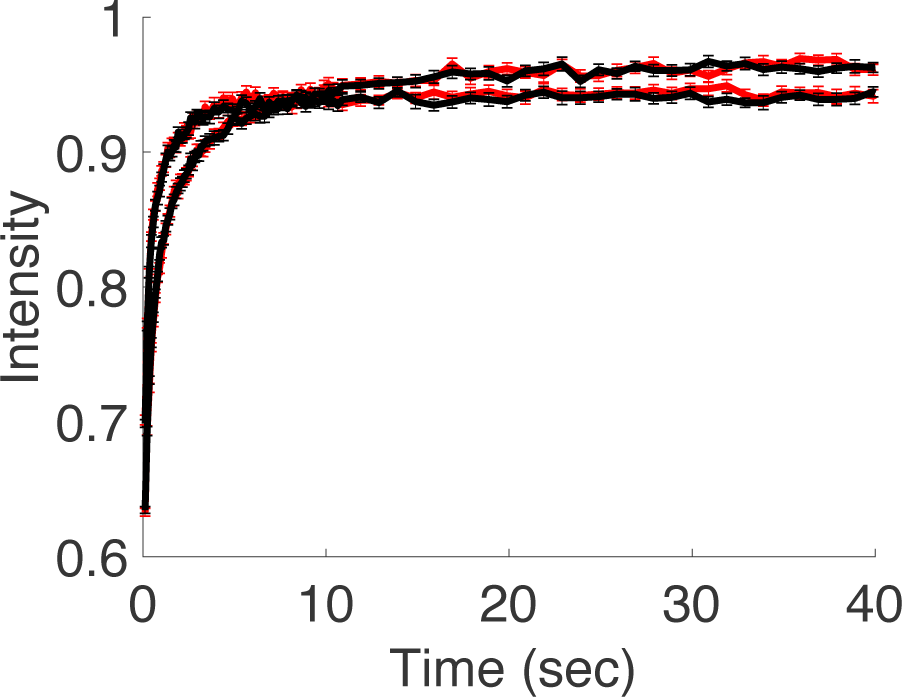
Validation of choice of simulation maximum time step. DCMS simulations for the moss geometry at the cell center and edge were performed with a maximum time step of 0.012 *s* (red) and 0.00012 *s* (black), respectively. *D* = 8.25 *μm*^2^s^−1^, n=50, error bars indicate standard error.

#### S1.3 Confocal Imaging and Bleaching

In order to accurately measure diffusion of an experimental fluorophore, it is important to model the imaging and photobleaching events on the confocal microscope properly[4]. Conventionally the Point Spread Function (PSF) of a confocal microscope is the result of the convolution of the excitation beam which can described by a Gaussian beam, i.e. a beam with a Gaussian profile along the *x* and *y* axes, and Lorentzian along the *z*-axis, and an emission beam which is typically modeled by an Airy function [5, 6]. While more detailed analytical calculations for the PSF of a confocal microscope exist [7], the required numerical integration would put a prohibitive load on the computational modeling when rendering images. To find the best parameters to empirically and efficiently describe the PSF of our microscope, we imaged 175 *nm* fluorescent beads as described in **Section 2.5**. As the beads used to measure this PSF are not an ideal point source, this produced a convolution of the PSF and the bead volume. After using a three-dimensional fitting routine in which a candidate PSF is convolved against a sphere representing the bead, we found that our experimental *z*-stacks were best fit by the convolution of two Gaussian beams (see **Fig. S3**). Thus, our resulting analytical approximation to the PSF takes the form

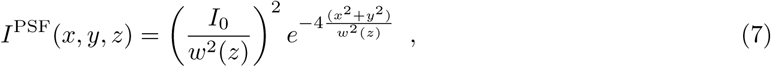

where the beam waist is given by

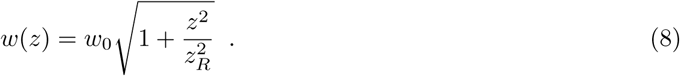

Here, *w*_0_ is the minimum beam waist and *z_R_* is the Rayleigh range. This expression resulted in an improved fit compared to the commonly used 3D Gaussian model—as evidenced by the sum of squares shown in **Fig. S3c**. The resulting parameters for the fitting routines to the squared Gaussian beam are shown in **Table S1** (top). This improvement is likely due to the large pinhole size used, 2 *AU* (Airy Units).

To illustrate that the fitting accuracy of the 3D Gaussian can be improved by decreasing the pinhole size, we measured the point spread function again for decreasing pinhole sizes **Fig. S4**. To collect more light and more accurately depict the PSF, we reduced the confocal scan speed to 400 *Hz*. As the pinhole size is reduced, the 3D Gaussian approximation improves, but never outperforms, the Gaussian beam approximation. A pinhole size of 2 *AU* was chosen for FRAP experiments to reduce out of focus light and collect enough light to measure an adequate fluorescence signal in moss. Although decreasing the pinhole size removes out of focus light, it also reduces the amount of light collected. At smaller pinhole sizes, such as those less than 1 *AU*, the laser power required to collect a sufficiently large fluorescence signal would result in increased photo-toxicity and acquisition photobleaching.

**Table S1.**
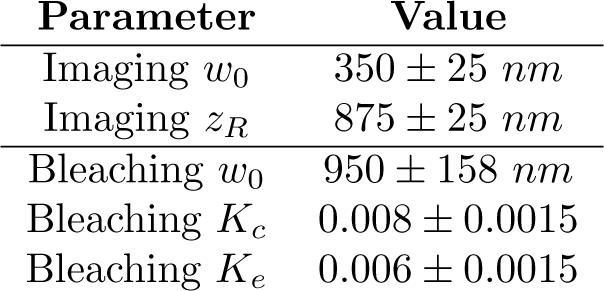
FRAP parameters determined for 3xmEGFP. Imaging parameters include the beam waist, *w*_0_ and Rayleigh range, *z_R_*. Confidence intervals are defined by the sampling distance for the respective parameter, as illustrated by the grid in **Figure S3**. Photobleaching parameters include the beam waist, *w*_0_, and bleaching probabilities, *K_e_* and *K_c_*, for the edge and center, respectively with 95% confidence intervals. *K*_*e*_ and *K_c_* Monte Carlo replicas never produced different parameters. For this reason the confidence intervals for these parameters are represented by our simulation sampling distance for the parameter *K*. It is important to note that using a shared bleach probability for the edge and center did not appreciably change our measured diffusion coefficient.

Since confocal scan time has been previously shown to influence fluorescence recovery [8], we performed a series of estimates to determine if modeling the entire scanning process was necessary. We estimated the distance a particle travels in a given time as the one-dimensional root-mean-square distance for a diffusive particle with diffusion coefficient *D* as 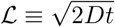. For *D* = 10 *μm*^2^*s*^−1^ (the largest value consistent with our experimental measurements in moss), we determined length scales for the durations of the various imaging processes, as shown in **Table S2**. We compared these distances with our imaging resolution—our PSF on the scale of hundreds of *nm*. The characteristic distance traveled during the acquisition of an image line is significantly smaller than the PSF size—defined by its beam waist, *w*_0_—while that of an entire image is larger. Thus, we chose a time step of 360 *μs* for imaging or photobleaching, a value equal to the time it takes to do a single line.

**Table S2.**
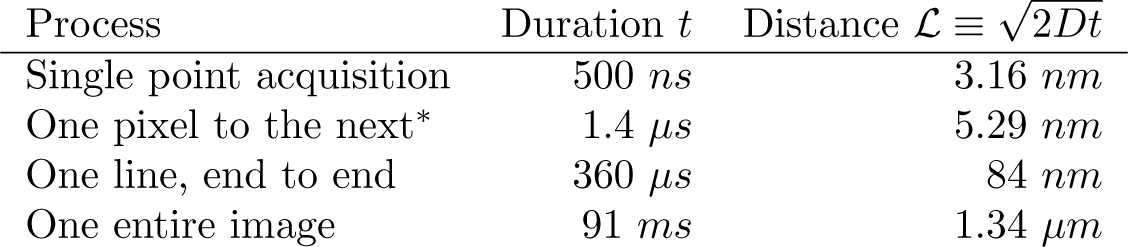
Estimates of the distance a particle with diffusion coefficient *D* = 10 *μm*^2^*s*^−1^ will travel during various processes of the Leica TCS SP5. * Total time, including both imaging and movement.

To simulate photobleaching on the confocal microscope, it was necessary to determine the spatial profile of an individual photobleaching event. This profile was characterized as a Gaussian beam, with a fluorophore-bleach probability (*P*_photobleach_) proportional to the beam intensity. As the microscope scans through a line at a speed much greater than diffusion (**Table S2**), we treated the photobleaching effect of a single horizontal line scan as a single instantaneous event. This was done by convolving the Gaussian beam intensity against a boxcar function from *x* = −*a* to *x* = *a*, i.e,

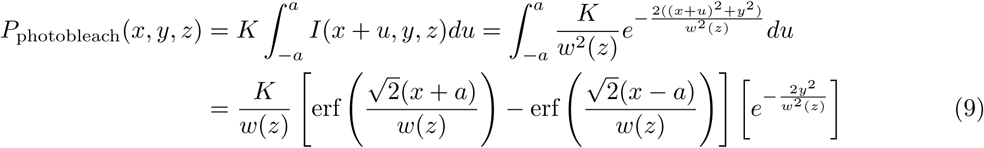

using the same *w*(*z*) functional form as used for imaging, with *K* as the bleaching proportionality constant. Simulated confocal scanning events using Eq. 9 are illustrated in **Fig. S5a and b**. To improve fitting to experimental data, we let the imaging beam width *w*_0_ differ from photobleaching width. The resulting parameters are shown in **Table S1** (bottom). It is not surprising that the photobleaching beam width differs from the imaging width because it has been previously shown that higher order diffraction rings can lead an extension of the bleaching width [9].

The entire FRAP process is illustrated in **Movie S1**, and **Movie S2**, with a snapshot from the simulations shown in **Fig. S2a**. Image stacks obtained from the simulations were analyzed to calculate the recovery curves, enabling the measurement of diffusion coefficients as explained in **Section S2.4**.

**Figure S3.**
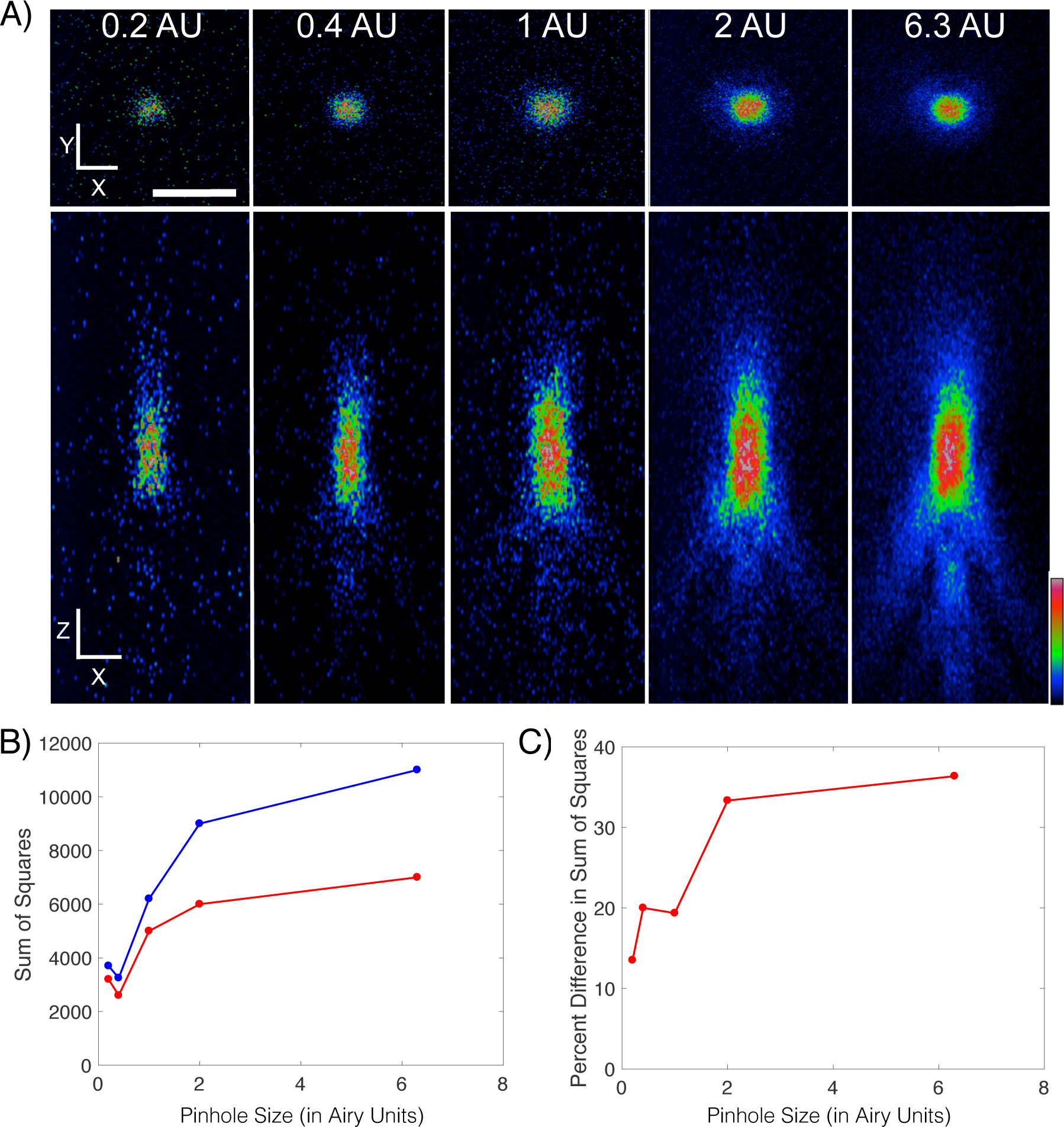
Measuring the imaging Point Spread Function (PSF). A) Squared Gaussian Beam approximation *I*^PSF^(*x, y, z*) (left, defined by Eq. 7) of experimental point spread function for 3xmEGFP (right). B) Parameter scan of potential values for *w*_0_ and *Z_R_*. Cool colors indicate small sum of squares between the experimental PSF and Squared Gaussian Beam, while warm colors indicate high sum of squares. C) Sum of squares between the experimental PSF and either the Squared Gaussian Beam (red) or the 3D Gaussian (blue).

**Figure S4.**
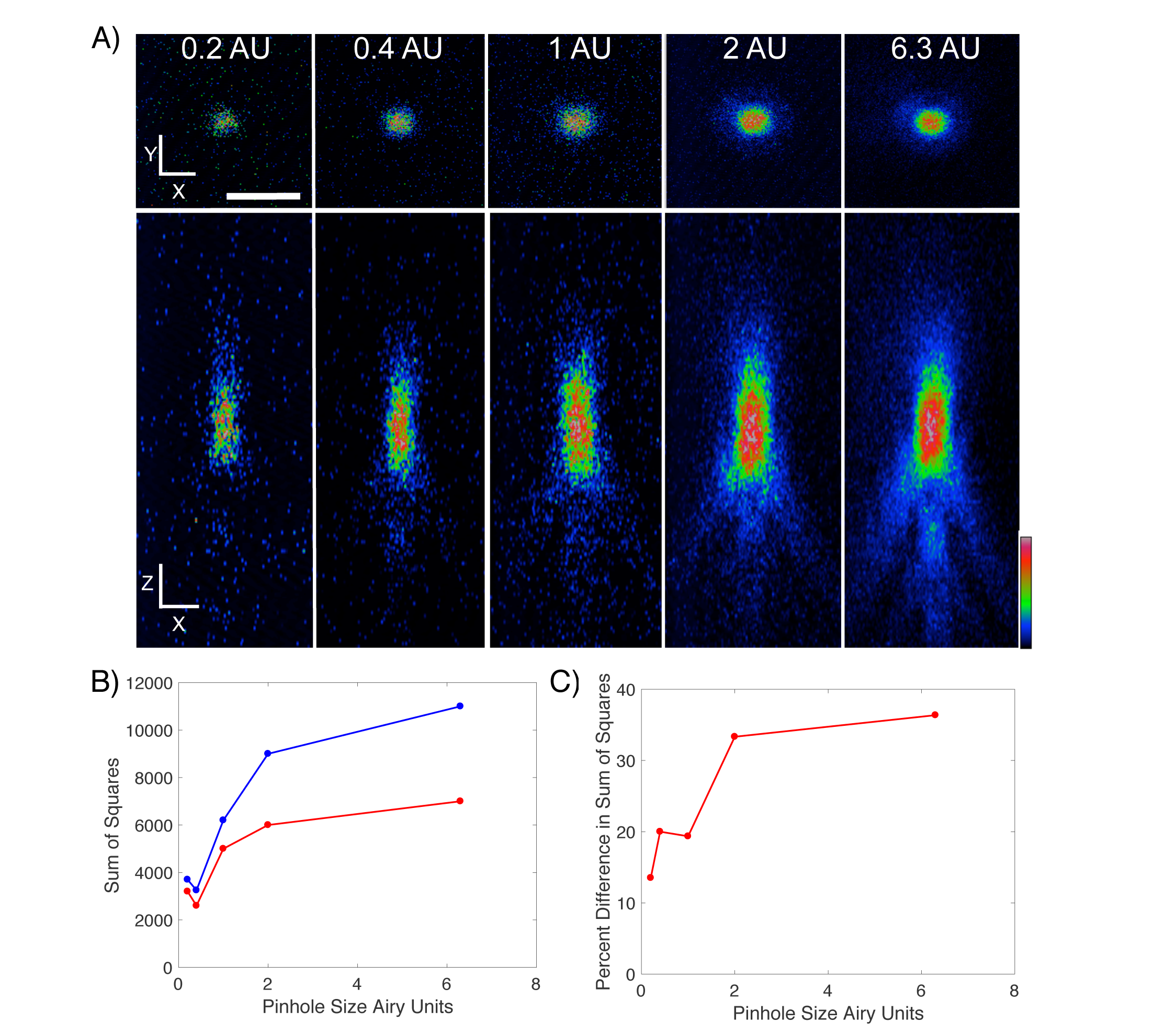
Effect of reducing pinhole size. A)Experimentally measured PSF in *x-y* and *x-z* for pinhole sizes of 0.2, 0.4, 1, 2, and 6.3 *AU*. Rainbow look up table indicates fluorescence intensity. Scale bar is 1 *μm*. B) Sum of squared differences for PSF fit with Gaussian beam (red) or 3D Gaussian (blue). C) Percent difference between sum of squared differences of the Gaussian beam and the 3D Gaussian approximations. Pinhole size is given in Airy Units (AU).

**Figure S5.**
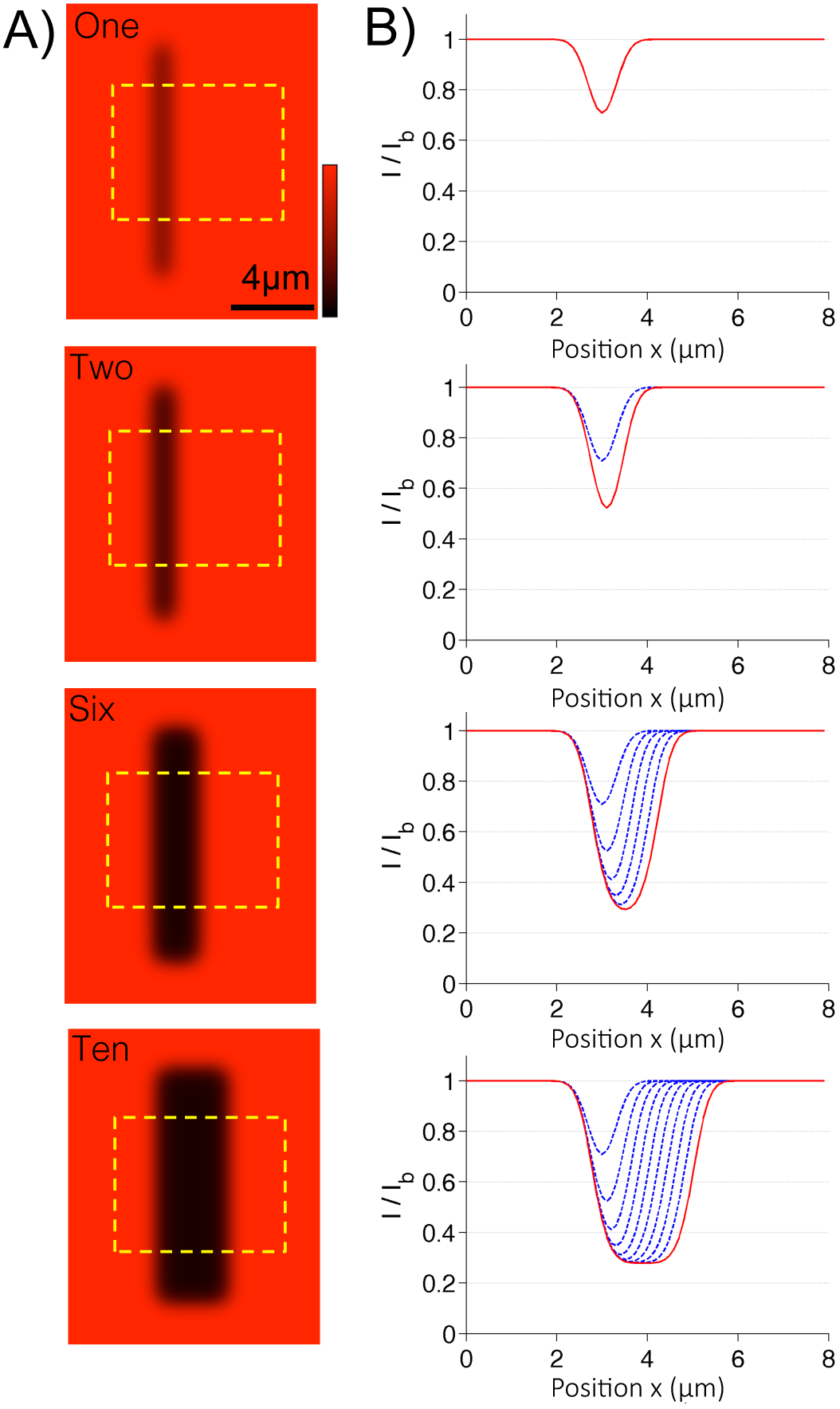
Modeling the photobleaching PSF A) Example simulated rectangular photobleaching with one, two, and ten confocal laser scans using Eq. 9. Dashed yellow rectangles indicate regions in which horizontal lines scans were measured and averaged along their vertical axis. B) Horizontal line scans through simulated photobleaching images in (A) for one, two, six, and ten confocal laser scans. *I* and *I_b_*, represent the simulated fluorescence intensity and the mean image background intensity, respectively. Blue dashed lines indicate the bleach result following each successive laser scan, and the red solid line indicates the final bleach result.

**Figure S6.**
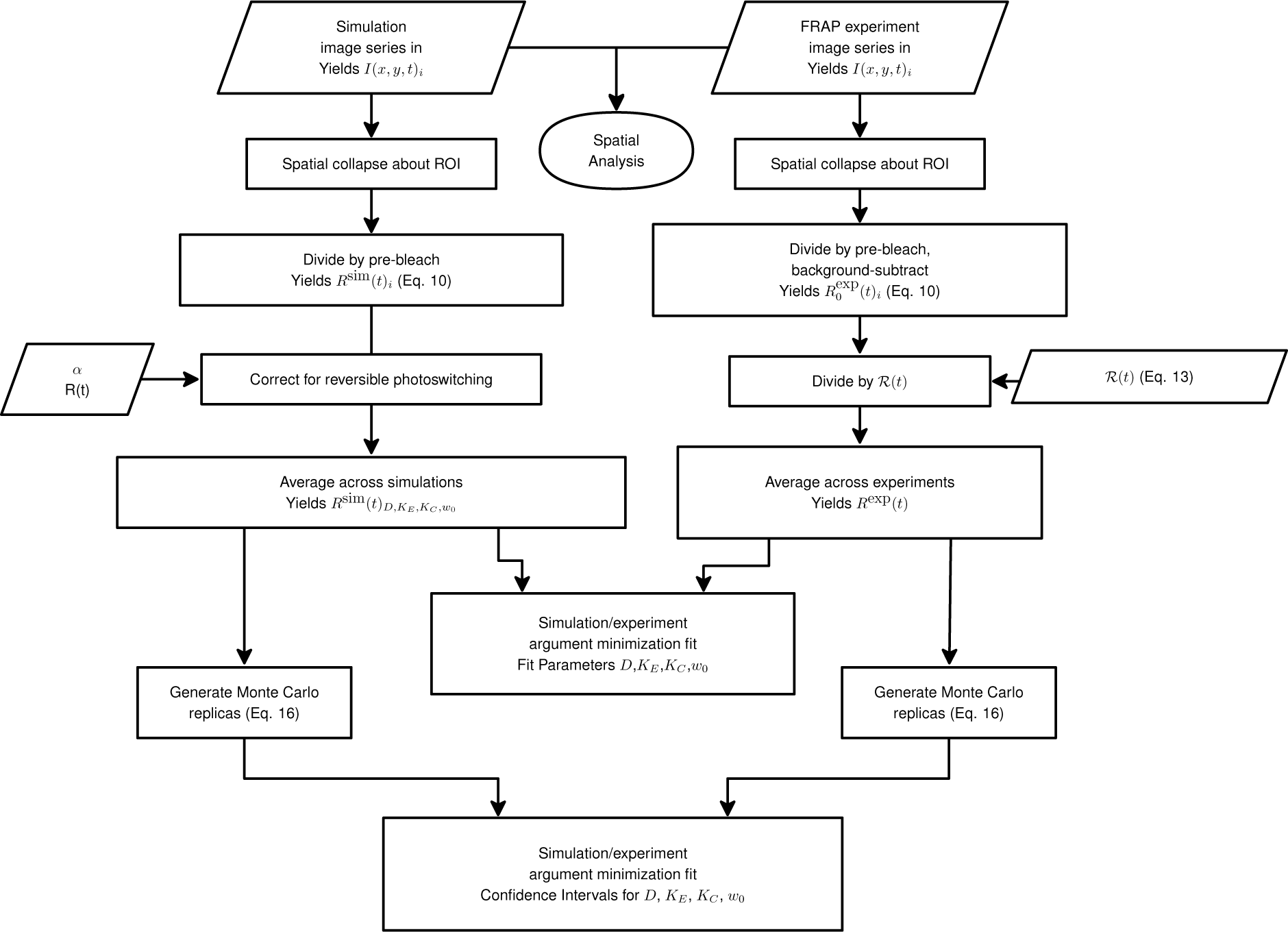
Flowchart for postprocessing of experimental and simulated images produced during FRAP. Both experimental and simulated images were reduced to recovery curves, averaged, and normalized as specified. The experimental curves were then subjected to an argument minimization to find their corresponding best fit simulations. Confidence intervals for the best fit parameters were generated via Monte Carlo replicas (**Section S2.5**).

### S2. Processing and Fitting of Experimental FRAP Data

#### S2.1 FRAP Post-Acquisition Image Processing

As depicted above, raw experimental FRAP tiff stacks *I*(*x, y, t*)_*i*_ were first analyzed with a custom ImageJ macro. The subscript *i* represents the *i^th^* replicate tiff stack for a particular experimental condition. The macro was written to track the photobleached ROI throughout the fluorescence recovery. The macro conducted background subtraction and divided the ROI fluorescence intensities by the ROI prebleach intensity to yield 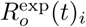 [10], namely,

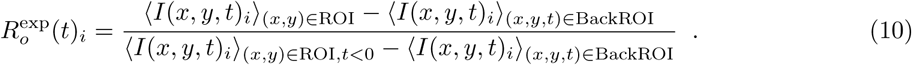

Here 〈…〉 denotes averaging about the corresponding subscripts inside the parenthesis, i.e.,

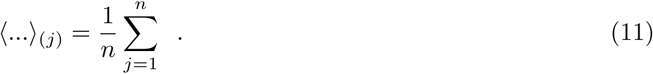

The subscript ROI represents the photobleaching ROI and the subscript BackROI represents the background ROI. All subsequent FRAP recovery curve processing was then conducted on the time dependent curves 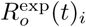. Simulated recovery curves were subjected to the same normalization without background subtraction to yield *R*^sim^(*t*)*_i_*.

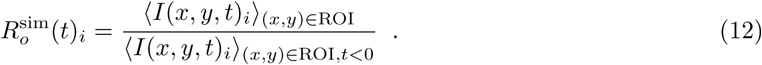

Experimental recoveries were then subjected to the acquisition photoblea.ching/reversible photoswitching correction (**Section S2.2**). Finally, using argument minimization, all relevant experimental parameters (**Section S2.4**) and corresponding confidence intervals were determined (**Section S2.5**).

#### S2.2 Image Acquisition Photobleaching and Reversible Photoswitching

Image acquisition photobleaching and reversible photoswitching during imaging, are processes by which fluorophores are photobleached or photo-converted by the laser used to excite the fluorophores. Depending on the experimental setup and conditions, such as the choice of fluorophore of interest and laser intensity, image acquisition photobleaching can significantly reduce fluorescence recoveries and lead to underestimation of the diffusion coefficient [11–13].

To measure the contribution of these effects, fluorescent cells were imaged with the same image sampling rate and experimental conditions as those used during FRAP experiments (**Section 2.2**). Cells were not subjected to the high intensity laser pulse necessary during FRAP experiments. Thus, reduced fluorescent intensity was primarily a result of the photobleaching and photoswitching during image acquisition. Images were then analyzed, and the mean fluorescence loss within the ROIs at the cell edge and center were measured across time. This ensured that the appropriate boundary effects were taken into consideration for each location. Fluorescence loss *R_L_*(*t*)*_i_* was divided by the intensity measured within the ROIs for the first image acquired *R_L_*(0)*_i_* to yield *R_Ln_*(*t*)*_i_*. Here, the subscript *i* denotes the *i_th_* experimental replicate. These normalized curves *R_Ln_*(*t*)*_i_* were averaged across these replicates < *R_Ln_*(*t*)*_i_* >_(*i*)_ to yield *R_Ln_*(*t*). Examples of these curves from the cell center of latrunculin B treated cells expressing 3xmEGFP are displayed in **Fig. S7a**.

**Figure S7.**
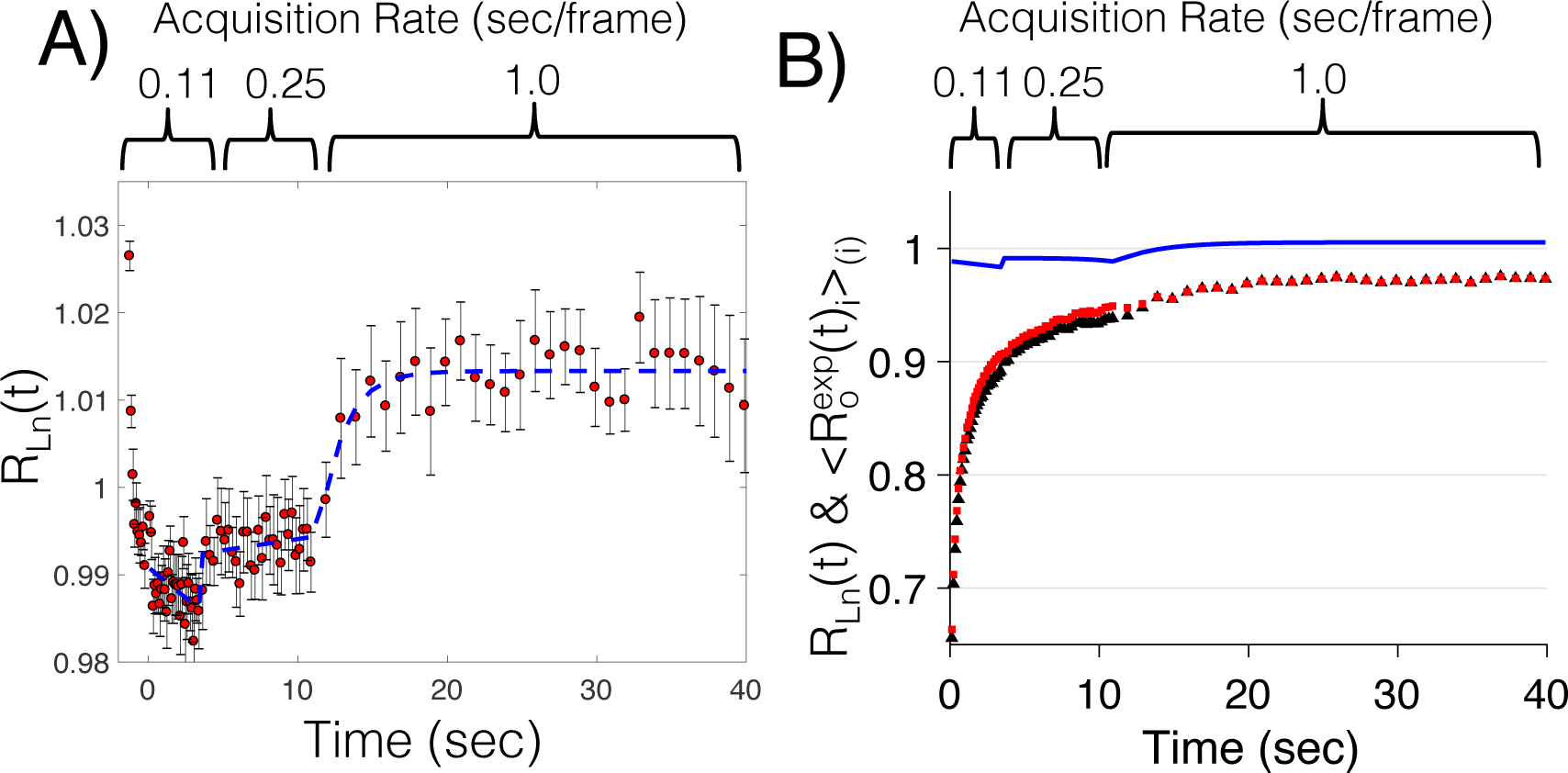
Image acquisition photobleaching and reversible photoswitching. A) Observed photobleaching and photoswitching during image acquisition at the center of cells expressing 3xmEGFP treated with latrunculin B. Red squares represent mean acquisition intensity, *R_Ln_*(*t*). Regions of different image sampling rates are outlined above. Blue dashed lines represents piecewise exponential approximation of image acquisition effects 𝓡(*t*), see Eq. 13. Error bars represent standard error. (n=9). B) Example fluorescence recovery before 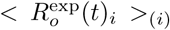 (black triangles) and after *R*^exp^(*t*) (red squares) correction for acquisition photobleaching and reversible photoswitching. Correction was conducted by dividing 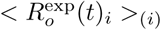 by the acquisition correction 𝓡(*t*) (blue) for the cell center of latrunculin B treated cells expressing 3xmEGFP. n=13 and 6, respectively.

The observed intensity profile shows a short bleaching event, followed by two longer recovery events at the input acquisition rates. This initial decay was less than 2% because we tried to minimize the laser power (10% power) required to illuminate our sample. Since our pinhole size was 2 Airy Units, we could collect sufficient light at low laser power 2.3. To help prevent further fluorescence loss, we then changed our sampling rate after the first 30 points were selected. This decrease in imaging sampling rate caused a small increase in fluorescence intensity, an effect which has previously been shown to be the product of reversible photoswitching during image acquisition [11–13]. Furthermore, this type of reversible photobleaching cannot be explained by a single light dependent rate constant that converts unbleached fluorophores to permanently bleached ones.

As a first approximation to this reversible photoswitching process, one would have to determine the rate constants that dictate the reversible bleaching reaction PB ← F ⇌ RB, where PB, F, and RB are the concentrations of permanently bleached, unbleached, and reversibly photoswitched fluorophores, respectively. However, this reaction can include additional states and transitions (such as from RB to PB), and the relevant rate constants have been shown to be light dependent [11], increasing the number of open parameters in the model that need to be experimentally determined. While DCMS is capable of reproducing this reversible photoswitching, to avoid a large reaction model and parameter scan, and increase the accuracy of our approach, we sought to remove acquisition photobleaching/reversible photoswitching effects from our experimental data directly [14].

Since these *R_Ln_*(*t*) curves were noisy, they were approximated by separate exponential functions as shown in **Fig. S7a**. Specifically, the three image sampling rates of 0.11, 0.25, and 1 s/frame used throughout the time course were each fit to their own exponential function, resulting in one curve, 𝓡(*t*), that approximates the entire image acquisition time course *R_Ln_*(*t*), i.e.,

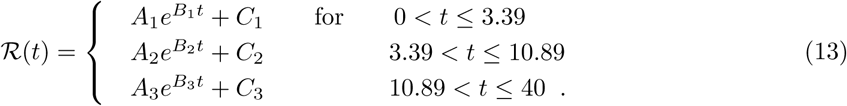

Finally, to correct for image acquisition photobleaching/reversible photoswitching each averaged FRAP experiment 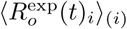 was divided by its corresponding acquisition approximation 𝓡(*t*) to yield *R*^exp^(*t*) as shown in the flow chart (**Fig. S6**). An outline of this correction procedure is provided in **Fig. S8**. Examples of fluorescent recovery curves before 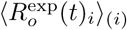 and after acquisition bleaching/reversible photoswitching correction *R*^exp^(*t*) can be found in **Fig. S7b**. Following correction, these recovery curves more accurately depict the dynamic nature of the fluorophores in question without being confounded by image acquisition photobleaching or reversible photoswitching, and can now be analyzed with a FRAP model that does not incorporate these effects.

**Figure S8.**
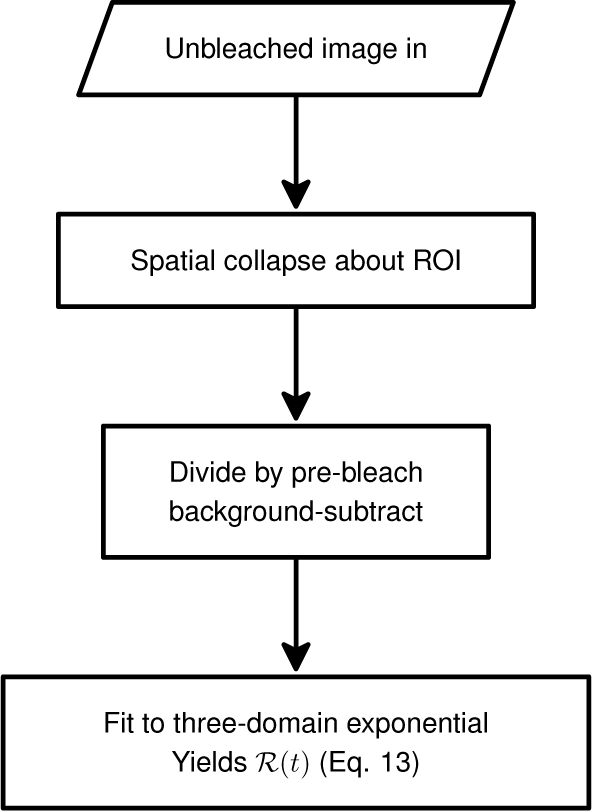
Flowchart for acquisition photobleaching/reversible photoswitching correction. Unbleached images were background subtracted, normalized by the prebleach fluorescence, and spatially collapsed about the ROI. These fluorescence traces were then fit to a three-domain exponential.

#### S2.3 Bleaching and Reversible Photoswitching

Similar to reversible photoswitching effects during acquisition, one should expect the main photobleaching event to be influenced by the photo conversion of the fluorescent protein [11, 12]. To measure the effects of reversible photobleaching, we performed photobleaching experiments in the nucleus of a cell line expressing nuclear GFP. During photobleaching, we illuminated the entire nucleus in plane, eliminating the effects of fluorescence recovery in the *x* and *y* directions. Based on the size of our PSF and our simulation evidence, fluorescence recovery in the *z* direction can be neglected. For these reasons, we attribute the fluorescence recovery seen in the nucleus to reversible photoswitching (see **Fig. S9a**). We found that the proportion of reversibly photoswitched molecules, *α*, was approximately 11%. To approximate the rate of reversible photoswitching, *R*(*t*), the fluorescence recovery was fit to a double exponential. As shown previously, [12], we found that *α* increased as we decreased the laser power of the main bleach event (**Fig. S9b**).

While DCMS can model this reversible photoswitching during bleaching, once again, for similar reasons, we opted to use an empirical approach and corrected for reversible photoswitching using a scheme devised by Mueller et al. [12]. Specifically, we modified all DCMS outputs with the correction scheme [12] expressed as

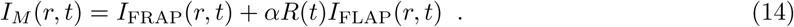

Here *I_M_* (*r, t*) represents the observed fluorescence recovery as a function of space and time for a photobleaching experiment with irreversible photoswitching, *I*_FRAP_ (*r, t*) represents the contribution of fluorescence recovery from unbleached fluorophores into the bleached region, and the expression *αR*(*t*)*I*_FLAP_ (*r, t*) is the contribution of fluorescence recovery due to reversibly photoswitched fluorophores. *I*_FLAP_ (*r, t*) is used here because the population of reversibly photoswitched fluorophores can be thought of as a Fluorescence Loss After Photobleaching (FLAP) experiment. FLAP is a process by which a high intensity laser pulse activates a population of fluorophores. The fluorescence intensity within this zone of activation will decrease as the activated fluorophores diffuse away. As shown by Mueller et al. [12], *I*_FLAP_(*r,t*) = [1 −*I*_FRAP_(*r, t*)] and [1 − *I*_FRAP_(*r, t*)] can be substituted into Eq. 14. This is true because the laser pulse in both FRAP and FLAP is the same. It then follows that the solution to the diffusion equation for FLAP is equal to one minus the FRAP solution. Since our simulation produces images with the intensity profile *I*_FRAP_ (*r, t*) including all the necessary spatial effects, *I*_FLAP_ (*r,t*) also includes spatial effects. Given our measured *R*(*t*) and *α*, we were able to correct all DCMS outputs for reversible photoswitching (including spatial effects) using this approach (**Fig. S9c** and **Fig. S9d**).

#### S2.4 FRAP Parameter Minimization

To identify the simulation parameters that best fit the recovery curves produced by experiments, a least squares minimization routine was used. In this routine, an exhaustive parameter sweep was simulated to generate a library of simulated recovery curves for all relevant parameter combinations. Then based on quality of fit to the experimental curves, the best fit parameters were identified. Specifically, averaged experimental recovery curves *R*^exp^(*t*) were characterized into classes 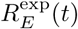 and 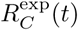, depending on if they were produced from the edge or center of the cell, respectively. Similarly, the averaged simulation recovery curves were characterized to produce 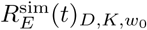 and 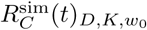. Here, *D* is the diffusion coefficient, *K* is the bleaching proportionality coefficient, and *w*_0_ is the bleaching probability beam waist used for the simulation. The best fit parameters for each experimental condition were found by minimizing the sum of squares differences between the experiment and simulation, for both the edge and center i.e.,

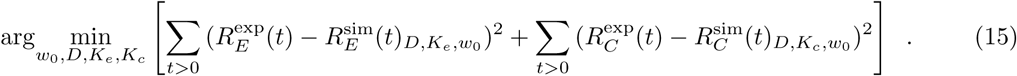

These parameters were found by allowing the edge and center to have the same diffusion coefficient *D*. The bleaching proportionality coefficient, *K*, was split into two independent parameters *K_e_* and *K_c_*, for the edge and center, respectively, to improve the resultant fit. The difference between *K_e_* and *K_c_* indicates that some underlying physical process may be altering the initial bleach depths at the edge and center (**Table S1**). Understanding the cause of this difference is beyond the scope of this paper. Note that this parameter minimization procedure was only applied to experimental data, using DCMS-generated recovery curves. Errors associated with the parameters were determined from the minimization procedure described in **Section S2.5**.

**Figure S9.**
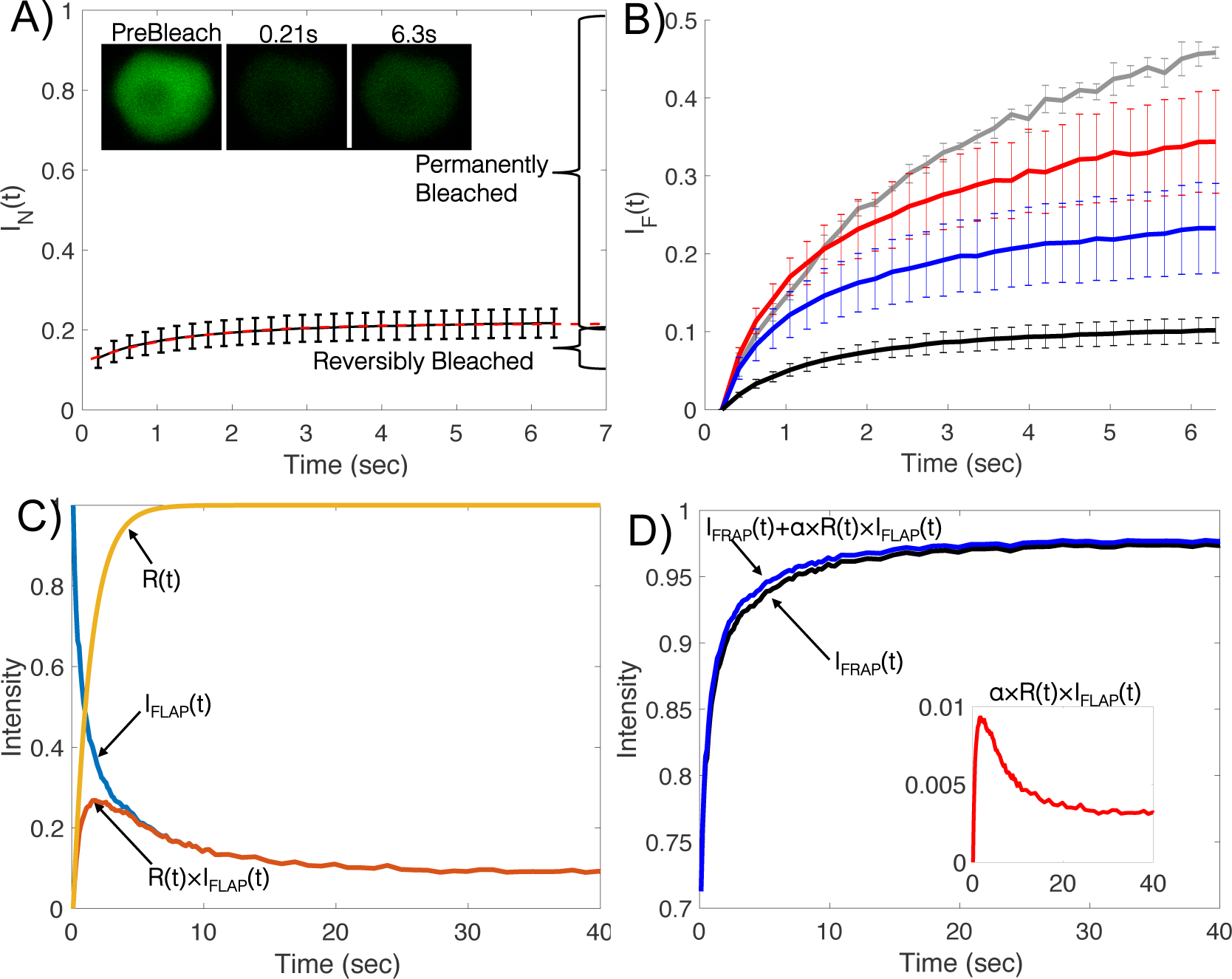
Reversible phot os witching caused by the main photobleaching event A) Measured reversible photoswitching of nuclear GFP, demonstrated by 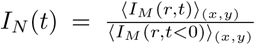. *α* denotes the percentage of reversibly switched fluorophores. n=4, error bars indicate standard error. Width of the nucleus images is 9 *μm*. B) Fluorescence recoveries, 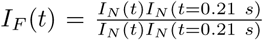, due to reversible photoswitching of nuclear GFP at 100% (black), 50% (blue), 30% (red), and 15% (grey) laser power. C) Illustration of reversible photoswitching components including the rate of reactivation *R*(*t*) (yellow), the rate of their movement out of the ROI *I*_FLAP_(*t*) (blue), and their product *R*(*t*)*I*_FLAP_(*t*) (red). D) DCMS-produced curves *FRAP*(*t*) (black) before correction for reversible photoswitching, the contribution of reversible photoswitching *αR*(*t*)*I*_FLAP_(*t*) (red inset), and the final corrected DCMS simulation *FRAP*(*t*) + *αR*(*t*)*I*_FLAP_(*t*) = *FRAP*(*t*) + *αR*(*t*)[1 − *I*_FRAP_(*r*, *t*)] (blue).

#### S2.5 FRAP Parameter Error Estimation

In order to provide a confidence interval on the parameters determined from the parameter minimization procedure, we used a Monte Carlo method. In the Monte Carlo method used, we took the mean and standard deviation for the average recovery curves, *R*^exp^(*t*) and *σ*^exp^(*t*), respectively, and generated a series of new recovery curves, *γ*(*t*)_*i*_, from them. This was done by sampling each point from a normal distribution with the parameters given by the experimental data, i.e.,

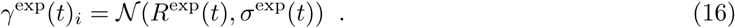

Here *i* represents the *i^th^* generated recovery curve. This generation process was repeated *n* times, where *n* matches that of the experiment, and averaged to produce *γ*^exp^(*t*). A similar procedure was then repeated on the simulation data (where *n* = 50), for all of the varying parameters, producing a new library of *γ*^sim^(*t*)_*D,K,w*_0__. The minimization procedure was then applied to find the *D, K_c_, K_e_*, and *w*_0_ for that set, as in Eq. 15,

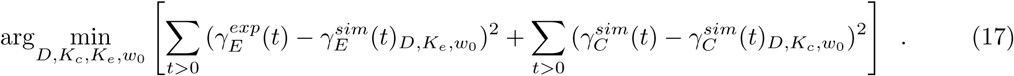

The random resampling imposed by both *γ^exp^*(*t*) and *γ^sim^*(*t*) allows for variation of the minimized parameters *D, K_c_, K_e_*, and *w*_0_ each time this routine is repeated. We then repeated this procedure until the standard deviation of the parameters *D, K_c_, K_e_*, and *w*_0_ reaches convergence. The confidence of these parameters was then expressed as two standard deviations of each parameter, as found in **Table S1**. Once again, this procedure was only applied to the experimental data, based on the DCMS-produced curves.

### S3. Spatial Analysis of Fluorescence Recovery

#### S3.1 Spatial ROI Extraction and Processing

To enable spatial analysis of fluorescence recoveries, experimental bleaching ROIs were extracted from image stacks with a custom cropping macro written in ImageJ. During cropping, the images were manually oriented and the macro was used to crop the ROIs from each image stack. Cropped image stacks were divided by the mean of their corresponding prebleach images and then averaged across replicate experiments, namely,

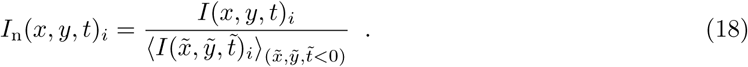

Here *I*(*x, y, t*)*_i_* is the *i^th^* image stack for a particular experimental condition, *t* < 0 denotes the prebleach portion of time in the image stack, and *I*_n_(*x, y, t*)*_i_* is the final normalized image stack. Simulated images were cropped and divided by the mean prebleach intensity in the same way.

To remove the effects of medial plane deviation and the limited accessible volume at the cell edge, described in **Section S5**, the extracted ROIs were again normalized. To conduct this second normalization, prebleach ROIs were averaged in time and along the vertical axis of the cell, as shown in **Fig. S10**. Zero valued pixels were ignored in this averaging. More specifically,

**Figure S10.**
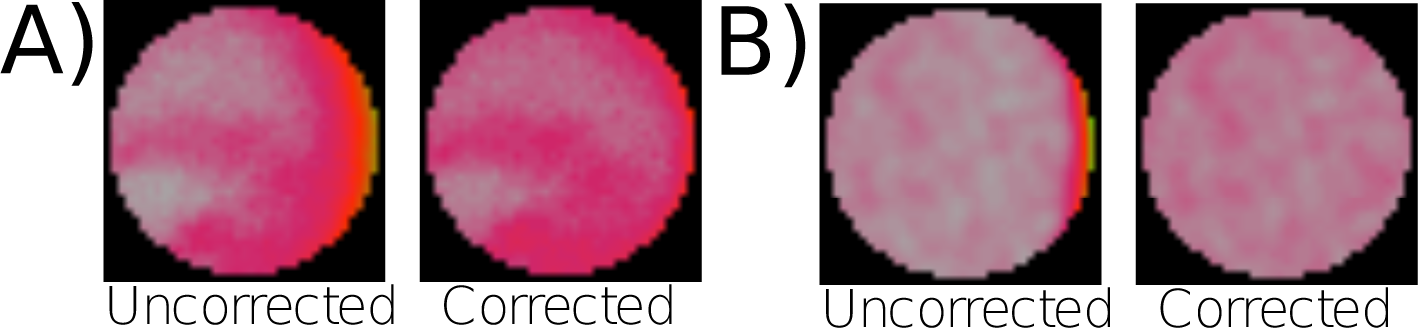
Accessible volume and medial plane deviation correction. A) 3xmEGFP at cell edge before (left) and after (right) accessible volume correction. (*n* = 13) B) 3xmEGFP simulation at cell edge before (left) and after (right) accessible volume correction. (*n* = 50) All cropped ROIs are 4 *μm* in diameter.

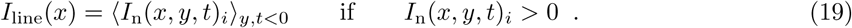

These line scans *I*_line_(*x*) were fit to an exponential function of the form *Ae^−Bx^* + *C*. Each ROI stack *I*_n_(*x, y, t*)*_i_* was then divided by this exponential decay to yield a volume corrected ROI *I*_v_(*x, y, t*)*_i_*, as shown in Fig. S10, i.e.,

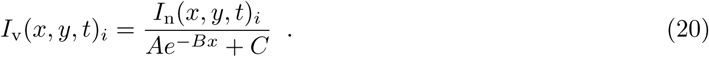

Volume corrected ROIs were used in all subsequent spatial analyses. An outline of this volume correction is depicted in Fig. S12.

#### S3.2 Fourier Analysis

Quantification of the directionality of fluorescence recovery was performed by subsampling the cropped ROIs into angular sectors about their centers. To reduce the noise associated with small regions of interest, a sliding window average was used. This sliding window was rotated 360 degrees and averaged about the horizontal axis of the ROI, producing an intensity profile that was dependent on the angular position of the sliding window, as shown in **Fig. S11**. To investigate fundamental modes of spatial recovery, the angular intensity profiles, *I*_v_, were reconstructed using a Fourier cosine series [15], namely,

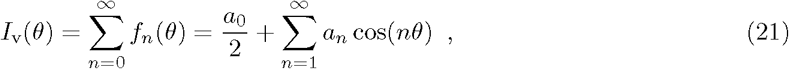

with *a_n_* denoting the amplitude of the *n*^th^ Fourier mode.

To illustrate the trends captured by the Fourier coefficients, three artificial gradients were generated and a Fourier analysis was used. The first gradient (**Fig. S11a**) increases linearly from right to left, while the second (**Fig. S11b**) peaks at 120 degrees. Finally, the third gradient (**Fig. S11c**) increases from right to left. The two linear gradients (**Figs. Slla** and **Sllc**) were well approximated by the first cosine mode of the Fourier series. This first (*n* = 1) amplitude describes the direction and steepness of the gradient. Specifically, a positive amplitude represents a gradient increasing from right to left, and a negative amplitude represents a gradient increasing from left to right. The mode amplitude value represents the steepness of the gradient. The gradient with a peak (**Fig. S11b**) requires the first three Fourier modes to be best captured. These simple examples illustrate how signs and amplitudes of the first few terms of the cosine series throughout fluorescence recovery make it possible to quantitatively characterize the directionality of spatial recovery.

**Figure S11.**
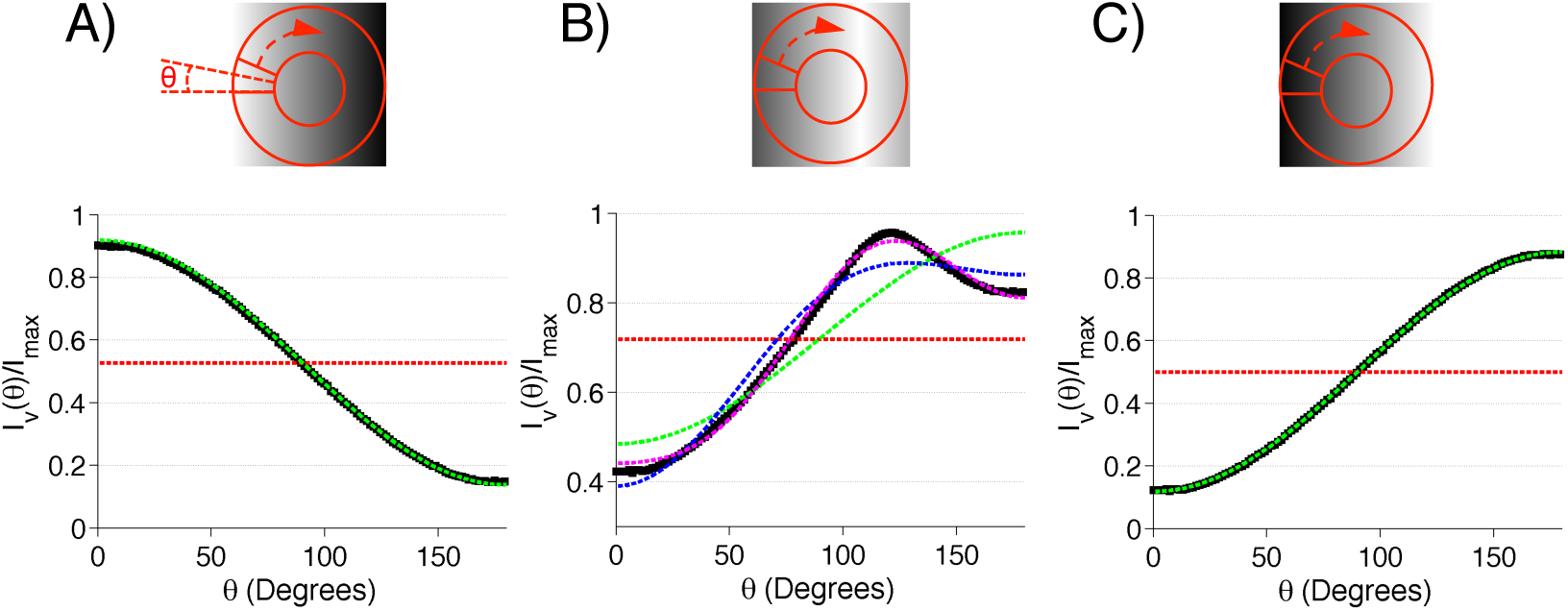
Fourier analysis of ROIs. A) Sliding angular intensity window for spatial analysis of generated gradients is indicated in red. Window is rotated 360 degrees and averaged about the ROIs horizontal axis. B) Fourier series representation of spatial intensity for generated gradients. Black squares represent the mean intensity of the sliding window, *I*_v_(*θ*), divided by the maximum pixel intensity of the image, *I*_max_. Dotted lines indicate *f_n_* where *f*_0_ is red, *f*_1_ is green, *f*_2_ is blue, and *f*_3_ is magenta (Eq. 21).

**Figure S12.**
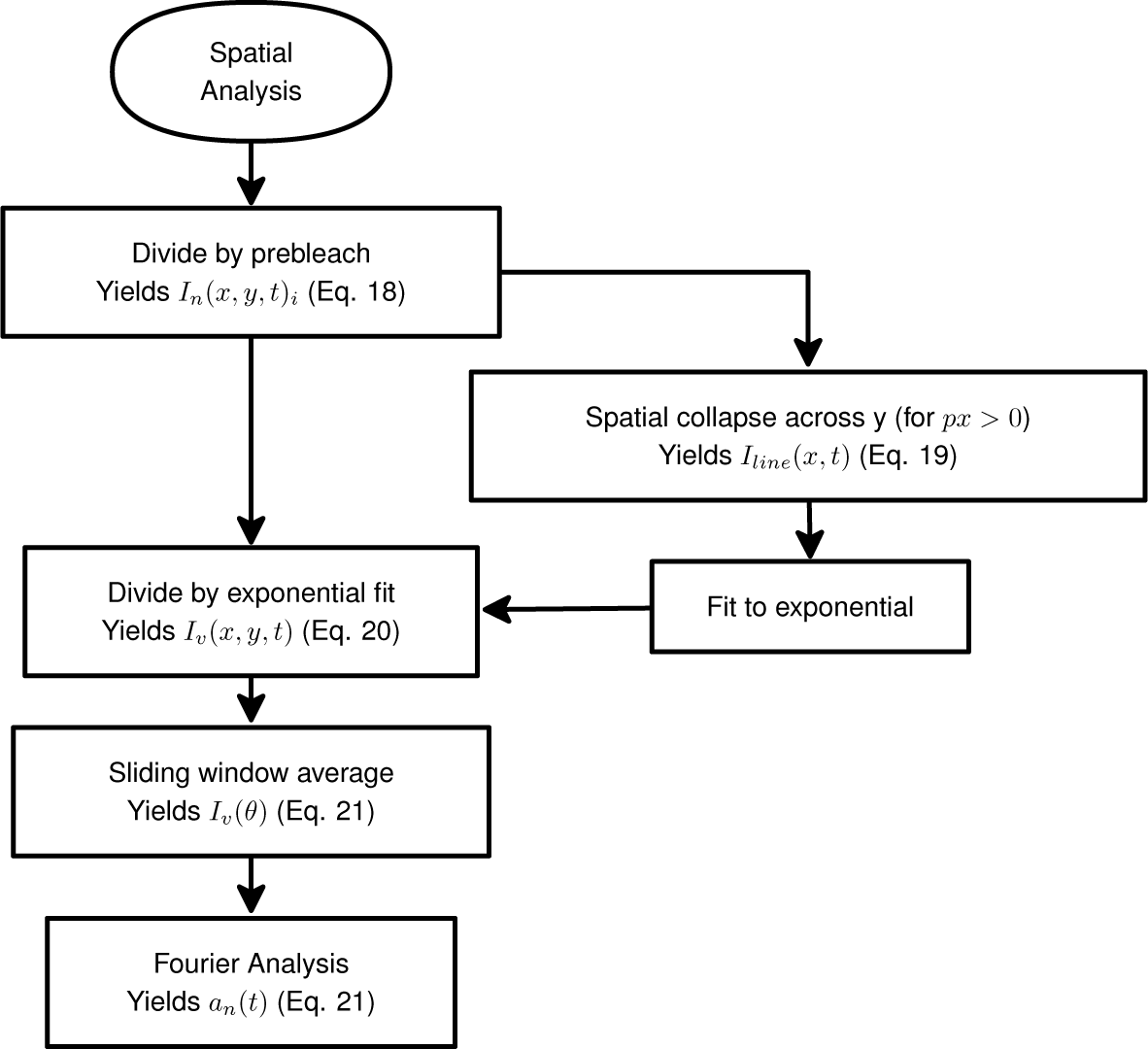
Flowchart of spatial analysis including volume correction and Fourier analysis. Image stacks prior to bleaching were subjected to line scans and fit to a single exponential. Post bleach images were divided by the exponential fit and quantified spatially with a sliding window average. Sliding window intensity profiles were analyzed with Fourier analysis.

#### S3.3 Fourier Analysis of 3xmEGFP at the Cell Edge

To characterize the directionality of fluorescence recovery, we cropped and averaged the photobleached regions (**Movie S3**). We could then detect intensity gradients as a function of angular position and time within the photobleached region (**Figs. S13a and S13b**). The shape of these gradients at early time points demonstrates that the geometry at the cell edge limited the direction of fluorescence recovery. The same analysis at the cell center yielded no observable gradient at early times. Additionally, the uniformity of fluorescence recovery at the cell center indicates that hydrodynamic effects do not influence 3xmEGFP dynamics (**Fig. S14**). This also indicates that the FRAP recovery at the edge is affected by its position next to the apical plasma membrane, such that material can not flow in or exchange in all directions as it can with the FRAP region in the center.

To quantitatively express the gradient’s transition from the transient state (Δ*t*_1_), found at early time points during recovery, to the steady state (Δ*t*_2_), and to test the statistical significance of the difference between the observed gradients, we used a Fourier cosine series to express the angular intensity profile at the cell edge (**Section S3.2**). Over the time course of the fluorescence recovery, the coefficients of the first fundamental mode of the Fourier series, *a*_1_(*t*) decayed to zero (**Fig. S13c**). At early times, Δ*t_1_*, the gradients exhibited time averaged Fourier coefficients significantly different than those observed during the steady state, Δ*t_2_*, (**Fig. S13d**). This significant difference between the time averaged coefficients at Δ*t_1_* and Δ*t_2_* indicates that experimental noise artifacts did not produce the gradient observed at Δ*t_1_*, and is consistent with a cell boundary effect on diffusion. The simulations at the cell edge recapitulated the angular intensity profiles observed transiently (Δ*t_1_*: 0 − 0.1*s*) and at steady state (Δ*t_2_*: 20 − 40 *s*) (**Figs. S13a-d** and **Movie S3**). This agreement between the simulated and experimental recovery supports the accuracy of our model of 3xmEGFP diffusion, and shows that spatial effects dominate over reversible photoswitching for this system.

**Figure S13.**
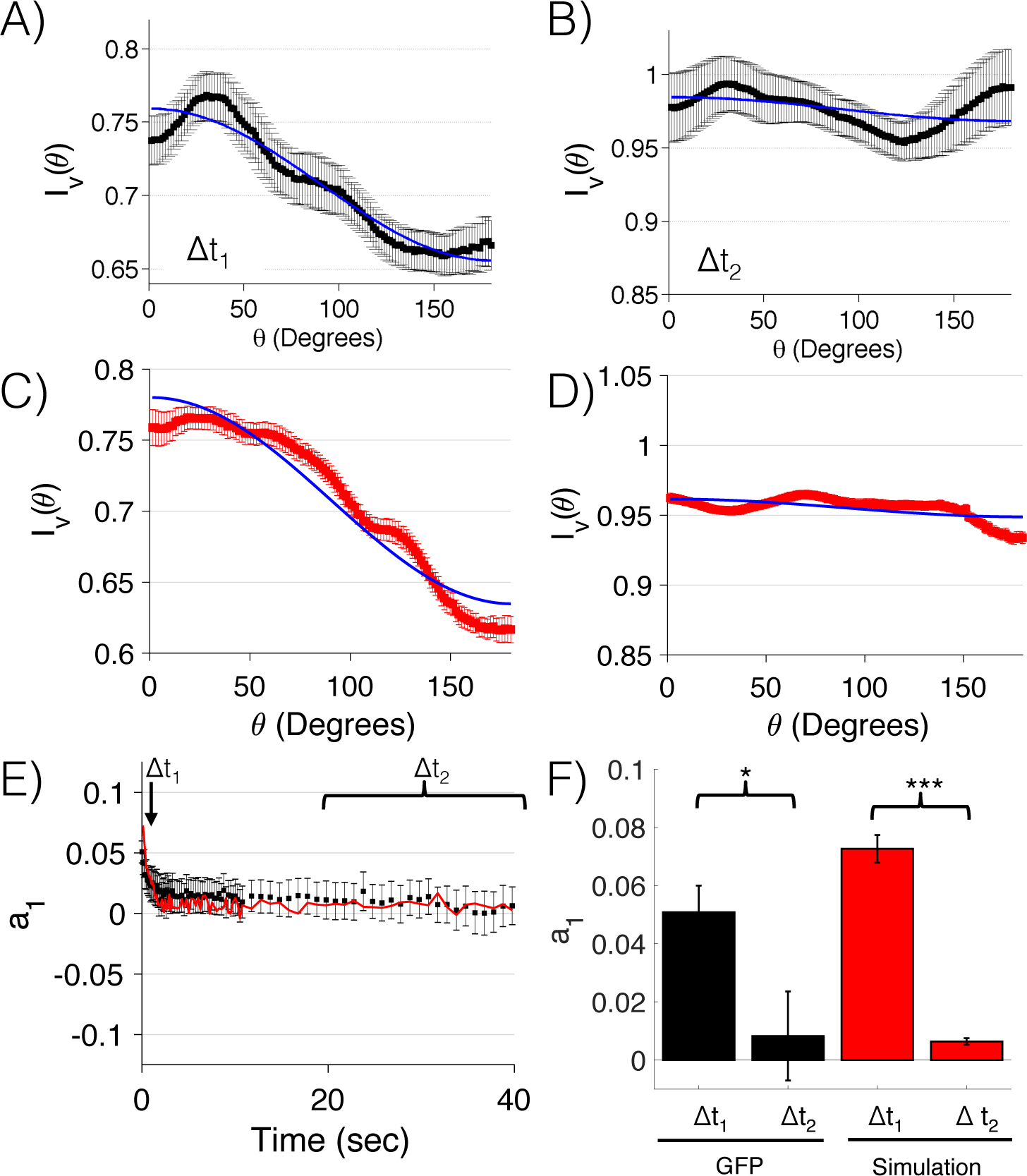
3xmEGFP Fourier analysis at the cell edge. A-B) Angular intensity profile of 3xmEGFP at the cell edge during the time intervals Δ*t*_1_ and Δ*t*_2_. Δ*t*_1_ 0 − 0.11 *s* (A) and Δ*t*_2_ = 20 − 40 *s* (B). First mode of the Fourier series, *f*_1_, is indicated in blue, *n* = 14, error bars indicate standard error. C-D) Angular intensity profiles of 3xmEGFP simulations at the cell edge during the same time intervals. Δ*t*_1_ = 0 − 0.1*s* and Δ*t*_2_ = 20 − 40 *s* (D). First mode of the Fourier series, *f*_1_, is indicated in blue. n=50, error bars indicate standard error. E) Amplitude of the first mode of Fourier series, ai, across time for 3xmEGFP (black) and simulation (red) for angular intensity profiles at the cell edge, *n* = 14 and 50 respectively; error bars indicate standard error. Δ*t*_1_ = 0.11 *s*; Δ*t*_2_ = 20 − 40 *s*. F) Fourier coefficients, ai, for frame averaged angular intensity profiles. 3xmEGFP cells are indicated in black and the simulations are indicated in red. Differences between Δ*t*_1_ and Δ*t*_2_ for simulation and 3xmEGFP are significant with, p-value < 0.05 and p-value < 0.001, respectively, *n* = 14 and 50, for experiment and simulation, respectively; error bars indicate standard error.

#### S3.4 Fourier Analysis of 3xmEGFP at the Cell Center

To verify that no hydrodynamic effects were present at the cell center, Fourier analysis was applied to the untreated 3xmEGFP FRAP experiments at the cell center. No significant horizontal recovery gradients were detected by analyzing the first mode of the Fourier analysis (**Figs. S14**), indicating that no cytoplasmic streaming currents were present at the cell center.

**Figure S14.**
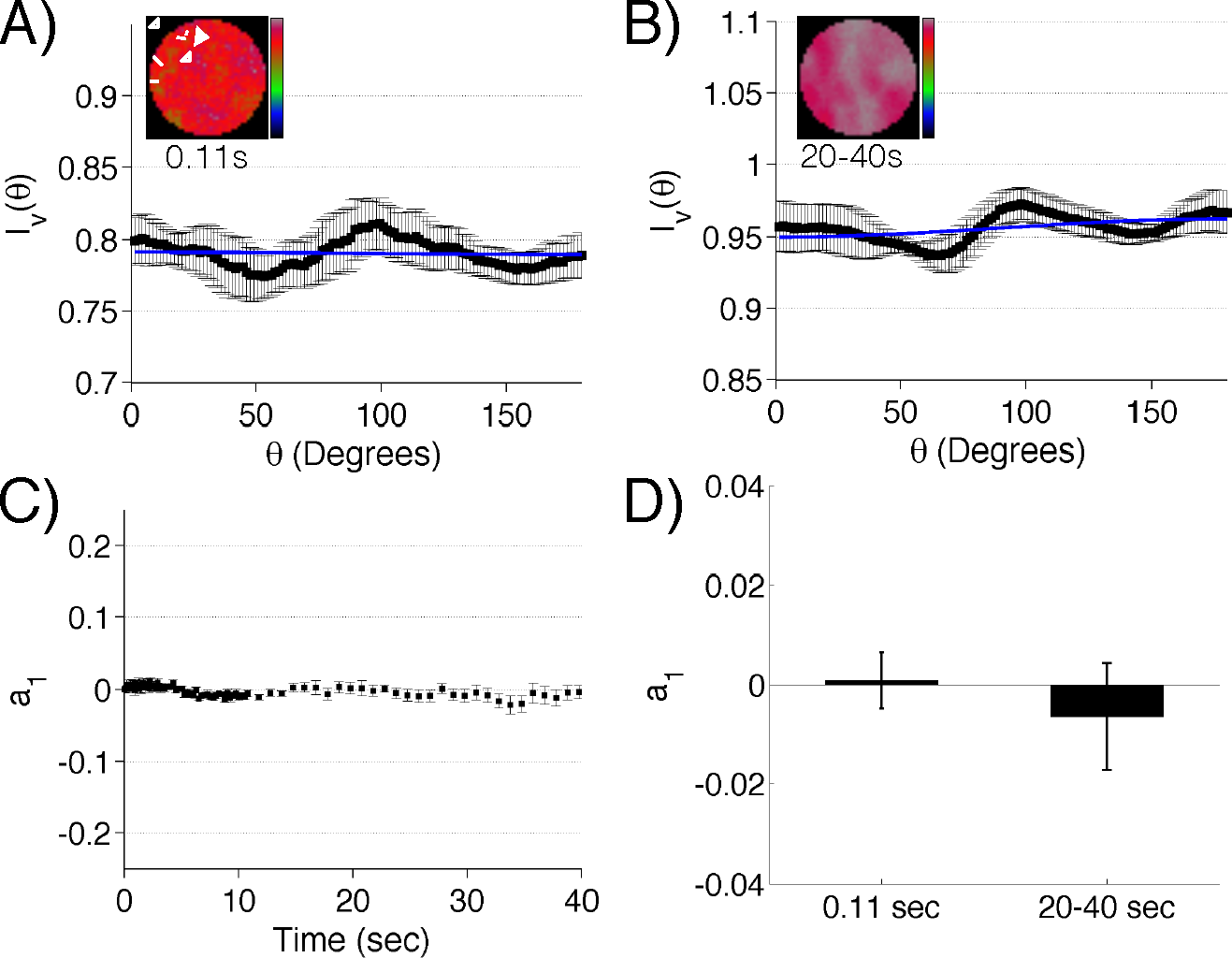
3xmEGFP center spatial analysis. A-B) 3xmEGFP spatial intensity profiles at 0.11 s (A) and at 20 − 40 *s* (B) following photobleaching. Black squares represent experimental intensity, and blue line indicates first mode *f*_1_ of the Fourier series. *I*_v_(*θ*) is the mean intensity within the sliding window for a particular angle *θ*. C) Amplitude of the first mode of Fourier series across time, *a*_1_. D) Binned first mode, *a*_1_, values at 0.11 *s* and 20 − 40 *s*. Error bars represent standard errors, p-value = 0.28 (n=7)

### S4. Detector Linearity

One of the main assumptions inherent in current analytical and computational FRAP models and analysis is detector linearity [16]. This assumption asserts that an increase in measured fluorescence intensity is linearly proportional to the increase of fluorescent particles within a region of interest, and holds well within a limited range, dependent on experimental setup. Due to the limited range of linearity of photon detectors in confocal systems, it was necessary to establish an operating range for the FRAP experiments in our experimental setup. Operating outside this linear range can produce results that are not comparable with those within the linear regime. To establish an operating range, a cell line with nuclear localized GFP was used. Repeated photobleaching was conducted to reduce the concentration of fluorescent particles within the nucleus, as shown in **Fig. S15**. FRAP recovery curves 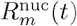 were generated by measuring the mean nuclear GFP intensity at each time point, *t*. Here, the subscript m denotes the *m^th^* successive bleach and the superscript *nuc* denotes that the intensity measured was from nuclear GFP. These curves were then divided by the nuclear GFP prebleach intensities 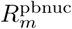 which corresponds to each successive bleach. By measuring fluorescence recovery at different concentrations we were able to determine if the concentration of particles could affect the resulting FRAP curves (**Fig. S15**). Since there was no difference in FRAP profiles across experimentally relevant intensities, it was concluded that these experiments were conducted within the linear operating range of the detector. Because the detector linearity assumption holds, no confocal detector corrections were applied to experimental FRAP data.

**Figure S15.**
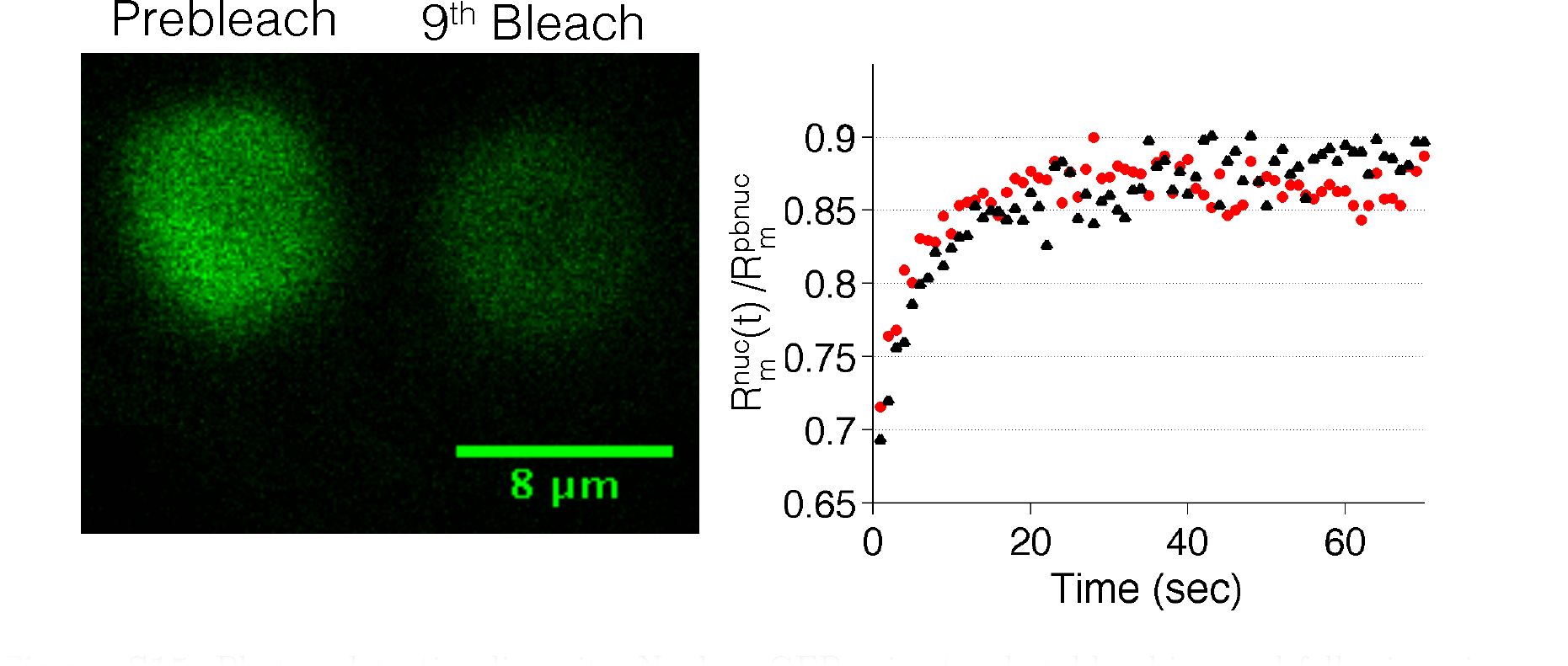
Photon detection linearity. Nuclear GFP prior to photobleaching and following nine successive photobleaching and recovery experiments (left). Example fluorescence recovery of nuclear GFP after the first (red circles) and ninth (black triangles) successive bleaching events (right). 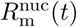 is the nuclear GFP fluorescent intensity and 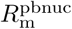 is the mean prebleach nuclear GFP intensity before each successive bleach.

### S5. Effects of Imaging and Photobleaching Away From the Medial Cell Plane

With and without treatment with latrunculin B, an intensity gradient was detectable at the edge of the cell for the 3xmEGFP fluorophores, **Fig. S16a** and **Fig. S10**. It is possible that these gradients are a result of imaging above and below the medial cell plane. These intensity gradients were observed prior to photobleaching and decayed toward the edge of the cell (**Fig. S16c**). Intensity gradients were found by taking line scans *L*(*x*)*_i_* through cells that were not subjected to photobleaching. Intensity gradients *L*(*x*)*_i_* were divided by their corresponding center intensity at *L*(0)*_i_* and averaged about the *i^th^* image, < *L*(*x*)*_i_*/*L*(0)*_i_* > _(*i*)_. Since these gradients were present in the 3xmEGFP cell lines, they were not the result of secretion at the cell edge. To determine if this reduced edge intensity could be a result of the reduced cell volume imaged at the edge [17], confocal images of uniformly distributed fluorophores were simulated, and the results for two different slices in *Z* are shown in **Fig. S16b**. In the medial plane at *Z* = 0, the reduced volume alone could not reproduce the experimental line scans. When imaged 2.5 *μm* above or below the medial cell plane, the simulations were able to produce a stronger gradient effect than the one observed experimentally, shown in **Fig. S16c**. This suggests that the experimental gradient is a result of imaging above and below the medial plane.

It is likely that the photobleaching experiments happened around the medial plane, *Z* = 0, with some variation in *Z* positioning plus or minus a few microns. To test the potential effects of this on rate of fluorescence recovery, FRAP experiments, with a diffusion coefficient of *D* = 0.35 *μm*^2^*s*^−1^ (to better resolve this effect), were simulated at the medial plane and 2 *μm* away from the medial plane, **Fig. S17a**. These simulated curves were averaged about each replicate 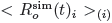 to yield *R*^sim^(*t*) for both imaging planes. Both recovery profiles for *Z* = 0 *μm* and *Z* = 2 *μm* shown in **Fig. S17b** are very similar. Moreover, randomly generated simulations with a 2 *μm* standard deviation away from the medial plane were generated, as displayed in **Fig. S17c**. These simulations had a constant diffusion coefficient of *D* = 0.35 *μm*^2^*s*^−1^ and were bound by recovery curves of simulations photobleached at the medial cell plane with diffusion coefficients *D* = 0.31 *μm*^2^*s*^−1^ and *D* = 0.39 *μm*^2^*s*^−1^. This error in diffusion coefficient measurement is well within a 15% error, as depicted in **Fig. S17c**.

Based on these two analyses, it can be concluded that imaging away from the medial plane can influence the spatial FRAP analysis, but photobleaching away from the medial plane has little influence on FRAP recovery curves. As a result, only spatial FRAP analysis was subjected to corrections to account for imaging away from the medial plane, as outlined in **Section S3.1**.

**Figure S16.**
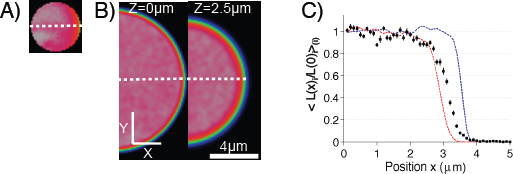
Intensity profile variation in Z. A) Experimental intensity gradient measured with unconjugated 3xmEGFP. Dashed white line indicates line scan through ROI at cell edge. ROI is 4 *μm* in diameter. B) Simulated intensity gradients for image acquisition medially at *Z* = 0 *μm* and 2.5 *μm*. above the medial plane. Dashed white lines indicate line scans through simulations. C) Horizontal line scans through simulation at *Z* = 0 *μm*. (dashed blue line) and *Z* = 2.5 *μm*. (dashed red line). Line scan through experimental 3xmEGFP intensity profile (black squares). < *L*(*x*)_*i*_/*L*(0)_*i*_ >_*i*_ denotes that the experimental line scan intensities were divided by the center intensity, and averaged across experimental replicates. Error bars indicate standard error. n=14 for experimental 3xmEGFP and n=50 for simulations.

**Figure S17.**
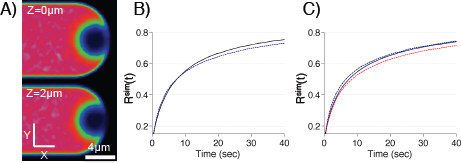
Recovery dependence on photobleaching plane position in Z. A) Simulated photobleaching medially at *Z* = 0 *μm*. (top) and 2 *μm*. above the medial plane (bottom). B) Simulated fluorescence recovery for photobleaching at *Z* = 0 *μm*. (dashed blue line) and *Z* = 2 *μm*. (black line). *R*^sim^(*t*) is the averaged simulation recovery curve. (n=50) C) Simulated averaged fluorescence recovery for randomly chosen FRAP experiments performed between the medial plane at *Z* = 0 *μm*. and 2 *μm*. above with a diffusion coefficient of *D* = 0.35 *μm*^2^*s*^−1^ (blue line). Dotted lines represent simulated medial fluorescence recovery at *Z* = 0 *μm*. with diffusion coefficients *D* = 0.31 *μm*^2^*s*^−1^ (dashed red line) and *D* = 0.39 *μm*^2^*s*^−1^ (dashed black line), respectively, and *R*^sim^(*t*) is the averaged simulation recovery curve. (n=50)

### S6. 3xmEGFP Latrunculin B Recovery

To verify that F-actin had little influence on the dynamics of freely diffusing species, cells expressing 3xmEGFP were treated with latrunculin B. Following latrunculin B treatment, the cells were subjected to FRAP at the edge and the center of the cell. Latrunculin B treatment had little effect on the recovery curves (**Fig. S18**) when compared to the corresponding untreated 3xmEGFP cells. Just as with untreated cells, latrunculin B treated cells displayed a reduced recovery at the edge when compared to the center. This trend was recapitulated by best fit simulations with a diffusion coefficient of *D* = 8.00±0.45 *μm*^2^*s*^−1^, which is within the error of our diffusion coefficient measured in untreated cells.

**Figure S18.**
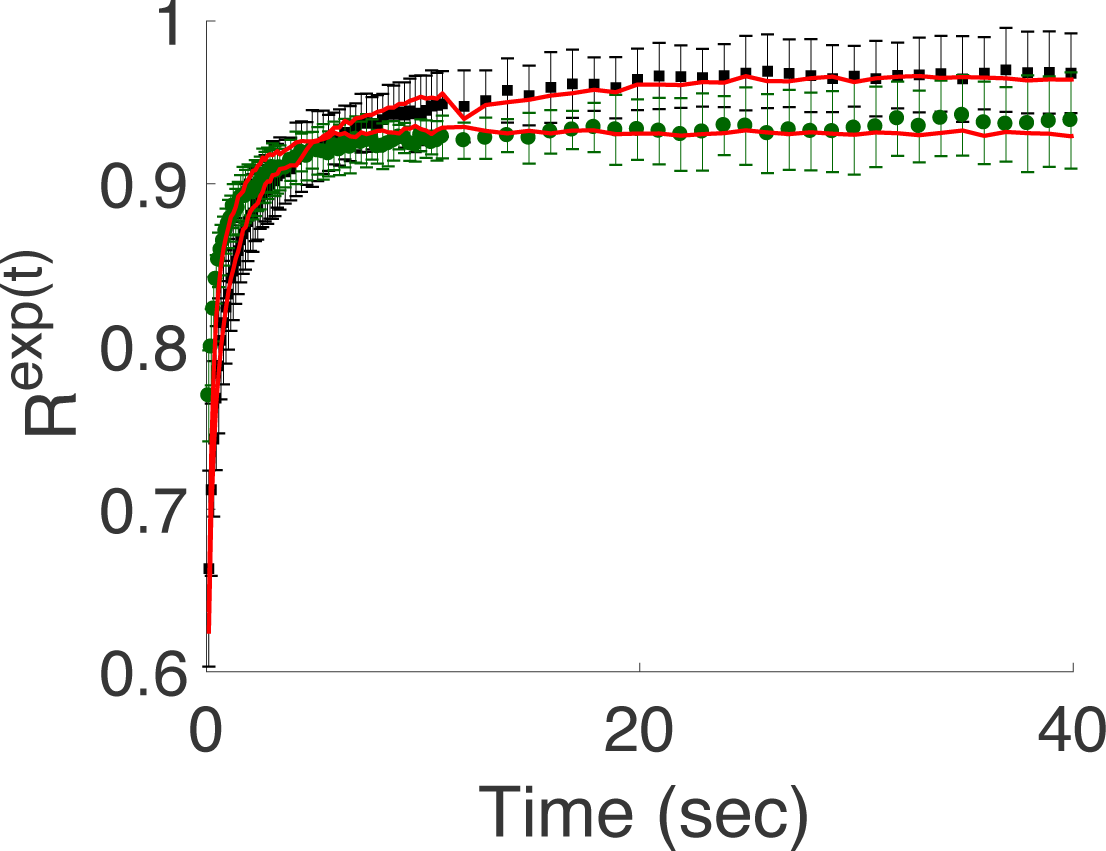
3xmEGFP latrunculin B FRAP. Fluorescence recovery of latrunculin B treated 3xmEGFP cells. Recovery curves, *R^exp^*(*t*), at the edge and center are indicated with black squares and green circles, respectively. *n* = 24 and 7 for the edge and center, respectively. Simulations corresponding to best fit parameters indicated in red where *D* = 8.00 ± 0.45 *μm*^2^*s*^−1^. Error bars represent standard deviation.

### S7. Analytical FRAP Models

In the following section, we briefly describe and provide functional forms for the analytical models we used for our fits. All models are normalized such that *FRAP*(0) = 0, and *FRAP*(∞) = 1. In order to fit to a recovery curve with bleach depth *I*(0) = *A*, and plateau *I*(∞) = *B*, these functional forms can be described with

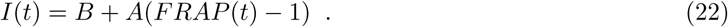

In what follows, *D* denotes the diffusion coefficient.

**Axelrod 1976b:** We use the second of the two models published in Axelrod’s paper introducing FRAP[18]. In this model, a Gaussian bleaching volume in a region with infinite boundary conditions is considered. Fluorescence recovery is given by

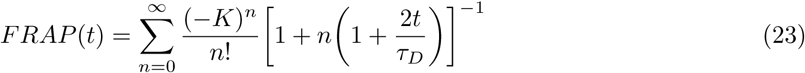

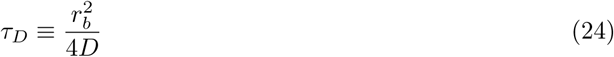

with the parameters

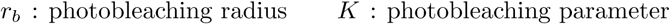

Note that the original paper additionally uses *P*_0_, the bleaching laser power, *A*, the light attenuation factor, and *C*_0_, the fluorophore concentration. We set these equal to unity to produce a normalization similar to the rest of the models presented here.

**Soumpasis 1983:** The Soumpasis model[19] uses similar assumptions to the Axelrod model, except using a sharp cylindrical bleach, rather than a two-dimensional Gaussian bleach. As its is a closed form expression using Bessel functions, it is very popular for curve fitting. Fluorescence recovery is given by

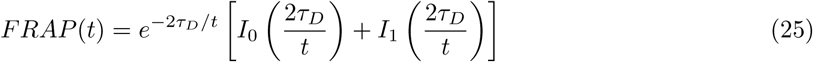

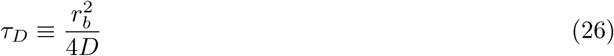

with the parameters

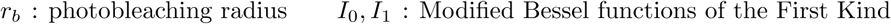

**Braeckmans 2003:** The Braeckmans model [20] incorporates the time required for the confocal to scan a large circular region of interest. It assumes that the region of interest is significantly larger than the imaging PSF, and uses a uniform cylindrical bleaching volume with infinite boundary conditions. Fluorescence recovery is given by

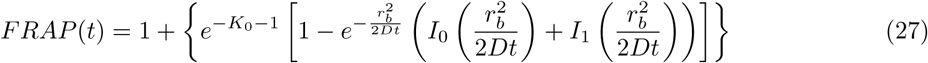

with the parameters

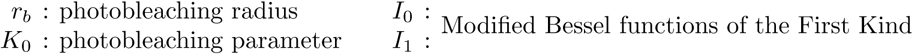

**1D FRAP Finite Boundary:** The 1D FRAP model[21] considers a ROI in the middle of a finite size strip with reflective boundary conditions at the ends. This is a very simple model, but is expected to work well for objects which are long and thin. Fluorescence recovery is given by

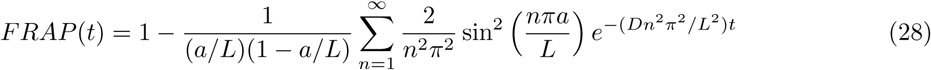

with the parameters

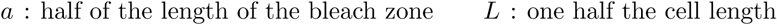

**1D FRAP Infinite Boundary:** The 1D FRAP considers a ROI in the middle of an infinite size strip. This is a very simple model, but is expected to work well for objects which are very long and thin. Detailed derivation of this model, and the subsequent 2D and 3D versions, are given in **Section S9**. Fluorescence recovery is given by

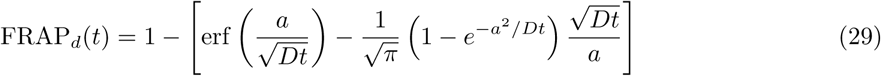

with the parameter

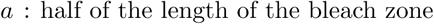

**2D Square FRAP** Infinite Boundary: The 2D FRAP considers a square ROI in the middle of an infinite 2D plane. Fluorescence recovery is given by

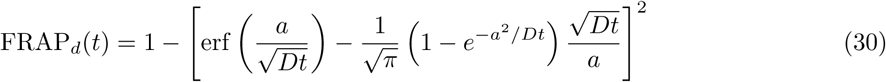

with the parameter

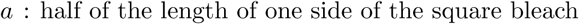

**3D Cubic FRAP in an Infinite Boundary:** The 3D FRAP considers a cubic ROI in the middle of an infinite 3D plane. Fluorescence recovery is given by

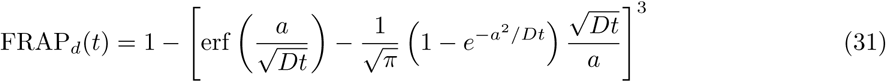

with the parameter

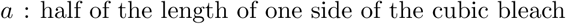

**Single Exponential (One Dynamic State):** The single exponential model represents a system with a single dynamic state. Fluorescence recovery is given by

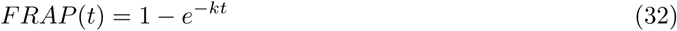

with the parameter

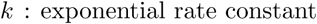

**Double Exponential (Two Dynamic States):** The double exponential model is often used to represent a system with two dynamic states. It is composed of the sum of two exponentials, each which represents a single state. Fluorescence recovery is given by

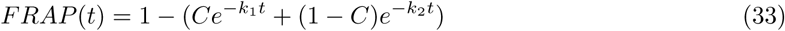

with the parameters

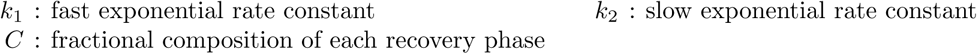

### S8. Analytical Demonstration of FRAP Boundary Effects in One Dimension

To better illustrate how a cell boundary can influence measured diffusion coefficients, we utilized the symmetry of the one-dimensional strip FRAP. For a cell of length 2*L*, and a bleach region of size 2*a* centered in the middle of the cell (as shown in **Fig. S19A**), the FRAP recovery curve is given by Eq. 28. If the bleach region is moved next to the boundary (as shown in **Fig. S19B**), the presence of the boundary slows down the fluorescence recovery. Due to symmetry, the bleach adjacent to the boundary (as shown in **Fig. S19B**) is equivalent to a centered bleach size of 4*a* in a cell of length 4*L* (as shown in **Fig. S19C**). Substitution of 2*L* and 2*a* into Eq. 28 shows that the expression containing the diffusion coefficient changes from (*Dn*^2^ *π*^2^/*L*^2^) to (*Dn*^2^ *π*^2^/*L*^2^). This clearly demonstrates that the recovery at the boundary (as shown in **Fig. S19B**) recovers like the centered solution (as shown in **Fig. S19A**) with an effective diffusion coefficient *D*_eff_ = *D*/4.

**Figure S19.**
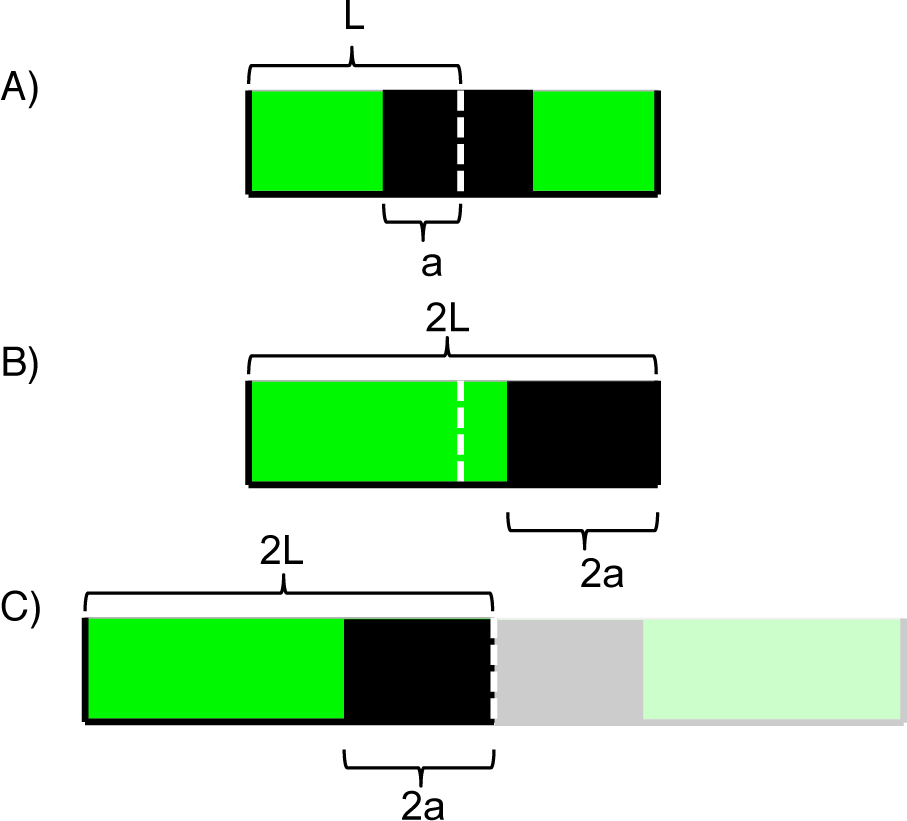
Different initial conditions for one-dimensional strip FRAP. A) A centered bleach region of length 2*a* in a cell of length 2*L*. B) A bleach region of length 2*a* at the right boundary in a cell of length 2*L*. C) A centered bleach region of length 4*a* in a cell of length 4*L*. The right hand side is tinted to illustrate the symmetry between the left hand side and (B). Fluorescence is indicated in green and bleach zone is indicated in black.

### S9. Asymptotic Behavior of FRAP Recovery

Solution of the diffusion equation in an infinite domain is known to exhibit power law scaling at long times. To further validate DCMS, we investigated the fluorescence recovery behavior at long times both analytically and using DCMS in one-, two-, and three-dimensions. While the analytical calculations are straightforward in Cartesian coordinates [22], we will briefly outline the steps here for completeness. Starting from the three-dimensional diffusion equation,

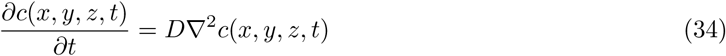

with the initial condition

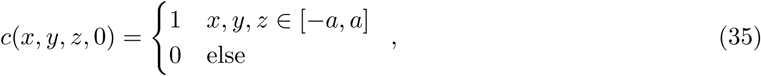

one can show that FRAP recovery is given by

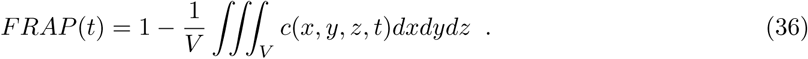

Using separation of variables, i.e. *c*(*x, y, z, t*) = *X*(*x*)*Y*(*y*)*Z*(*z*)*F*(*t*), and the diffusion equation (Eq. 34), one can show that the concentration at *t* = 0 can be written as

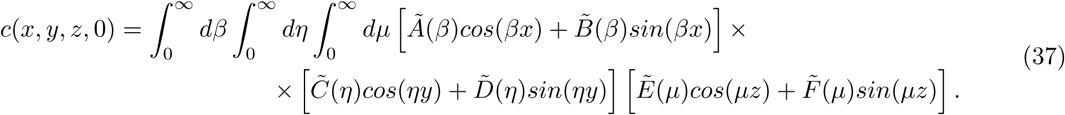

The coefficients 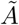 through 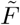, can be determined from the initial conditions using the orthogonality relations of the trigonometric functions. One can then show that the general solution can be written as [23]

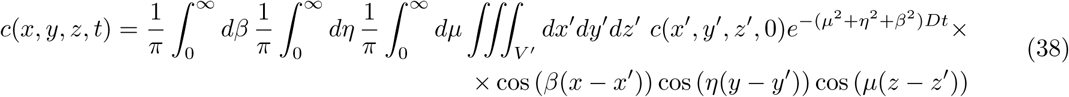

Integrating over *β*, *η*, and *μ* with the initial condition in Eq. 35 yields

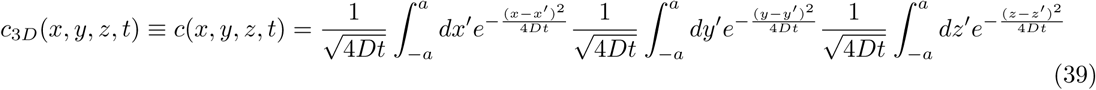

Performing the Gaussian integrals, one gets for the concentration profile in three-dimensions

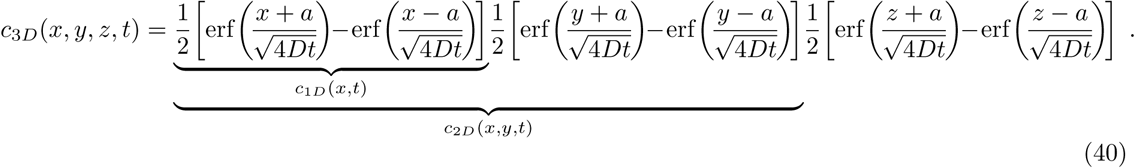

where *c*_1*D*_(*x, t*) and *c*_2*D*_(*x, y, t*) indicate the corresponding solutions in one- and two-dimensions. The FRAP recovery curve then can be calculated using Eq. 36 which yields its most general form

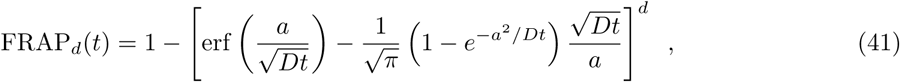

where *d* = 1, 2, 3 denotes the dimension. In the limit of long times, it can be shown that the asymptotic expansion is given by

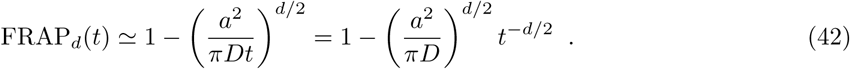

In order to validate DCMS with respect to this asymptotic behavior given by Eq. 42, we performed simulations within filopodia (*d* = 1, *t*^−1/2^), **Fig. S20A**, lamellipodia (*d* = 2, *t*^−1^), **Fig. S20B**, and a three-dimensional cube (*d* = 3, *t*^−3/2^) **Fig. S20C**. The results are in excellent agreement with theoretical predictions. In contrast, the single (Eq. 32) and double (Eq. 33) exponential models do not obey this scaling, once more illustrating the fact they do not have the right functional form (see **Fig. S20D**).

**Figure S20.**
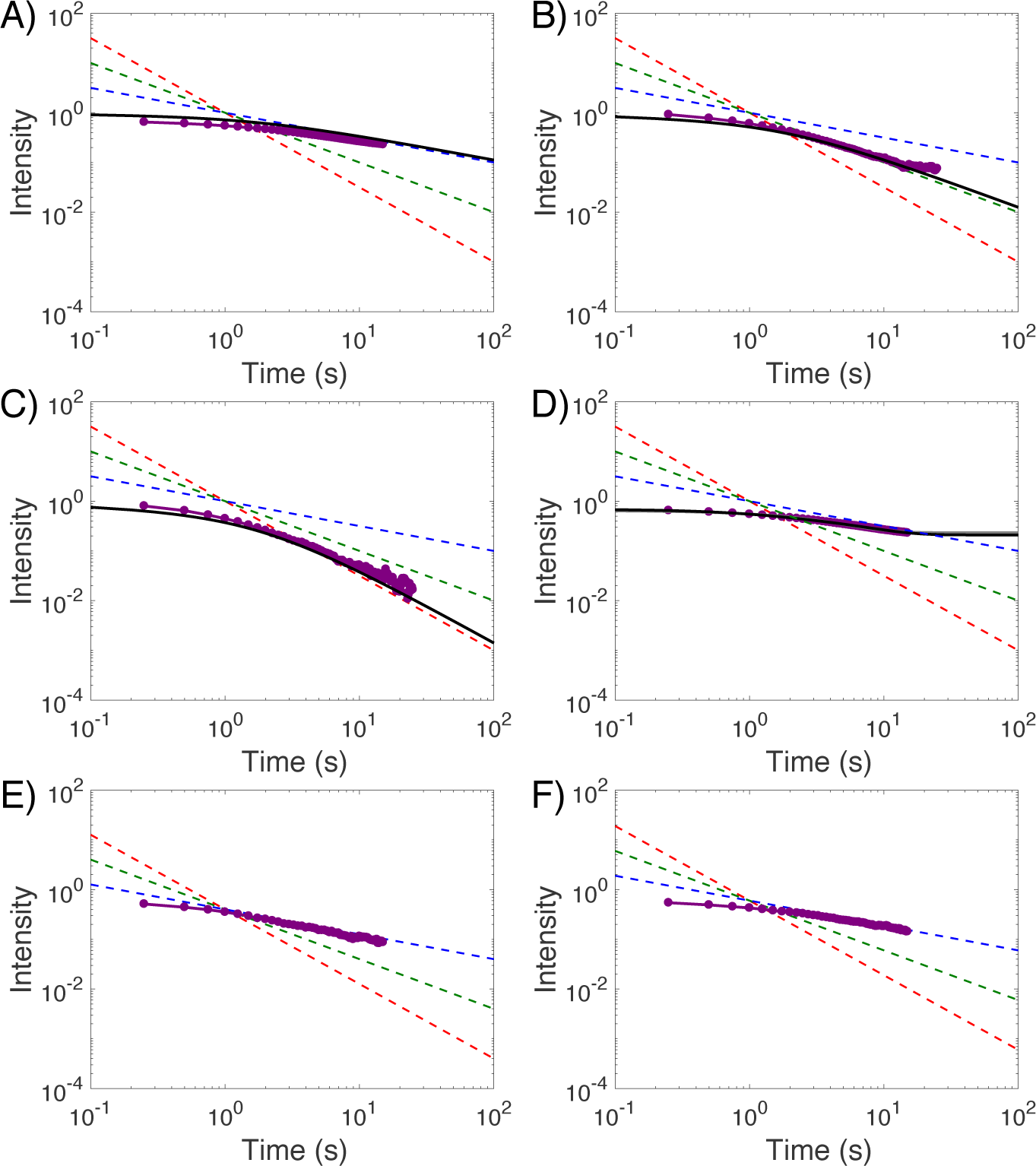
Asymptotic behavior of FRAP recovery illustrated using DCMS simulations. Red, green, and blue dotted lines represent Eq. 42 with *d* = 1, 2, and 3 producing *t*^−1/2^, *t*^−1^, and *t*^−3/2^ scaling, respectively. All fluorescence recovery curves were plotted as one minus the fluorescence intensity on a logarithmic scale. A) DCMS fluorescence recovery in filopodia with a centered bleach (purple) compared with Eq. 41 (*d* = 1, black line), showing *t*^−1/2^ scaling at long times. B) DCMS fluorescence recovery in lamellipodia with a centered square bleach (purple) compared with Eq. 41 (*d* = 2, black line), showing *t*^−1^ scaling at long times. C) DCMS fluorescence recovery in a large cubic cell with a cubic bleach (purple) compared with Eq. 41 (*d* = 3, black line), showing *t*^−3/2^ scaling at long times. D) Single (Eq. 32, gray line) and double (Eq. 33, black line) exponential fits to the filopodia center bleach (purple) do not exhibit power law scaling at long times. E-F) DCMS fluorescence recoveries (purple) in the moss geometry with bleaches at the cell center (E) and cell edge (F) exhibit long time power law behavior of the form *t*^−1/2^.

It is important to note that this scaling behavior can be affected in confined cellular geometries. For instance, simulations in the moss geometry with both a center and edge bleach exhibited power law behavior that converges to −1/2 instead of −3/2, as shown in Figs. S20E and F. This is not surprising since in the moss geometry, the combined effect of scanning that bleaches the ROI (see Fig. S21) and the fact that the moss cell has an aspect ratio like a tube, practically reduces this geometry to a onedimensional object in the limit of long times. However, at earlier times, this one-dimensional approach does not work, and the recovery is best explained by a two-dimensional model as illustrated in Table 1; since there is no z diffusion at early times, diffusion is practically in *x*– and *y*– directions.

### S10. FRAP Fitting Results for Various Cell Types

All model fitting to the simulations (except for the VCell VirtualFRAP tool) was performed in Matlab using the fitting function *lsqcurvefit* with equal weights for each point in the recovery. For VCell VirtualFRAP, we use the built-in function to produce a diffusion coefficient from an input image stack. This function was chosen because it enables the user to specify upper and lower bounds for parameter estimation. The results of these fits can be found in **Table I**.

**Table S3.**
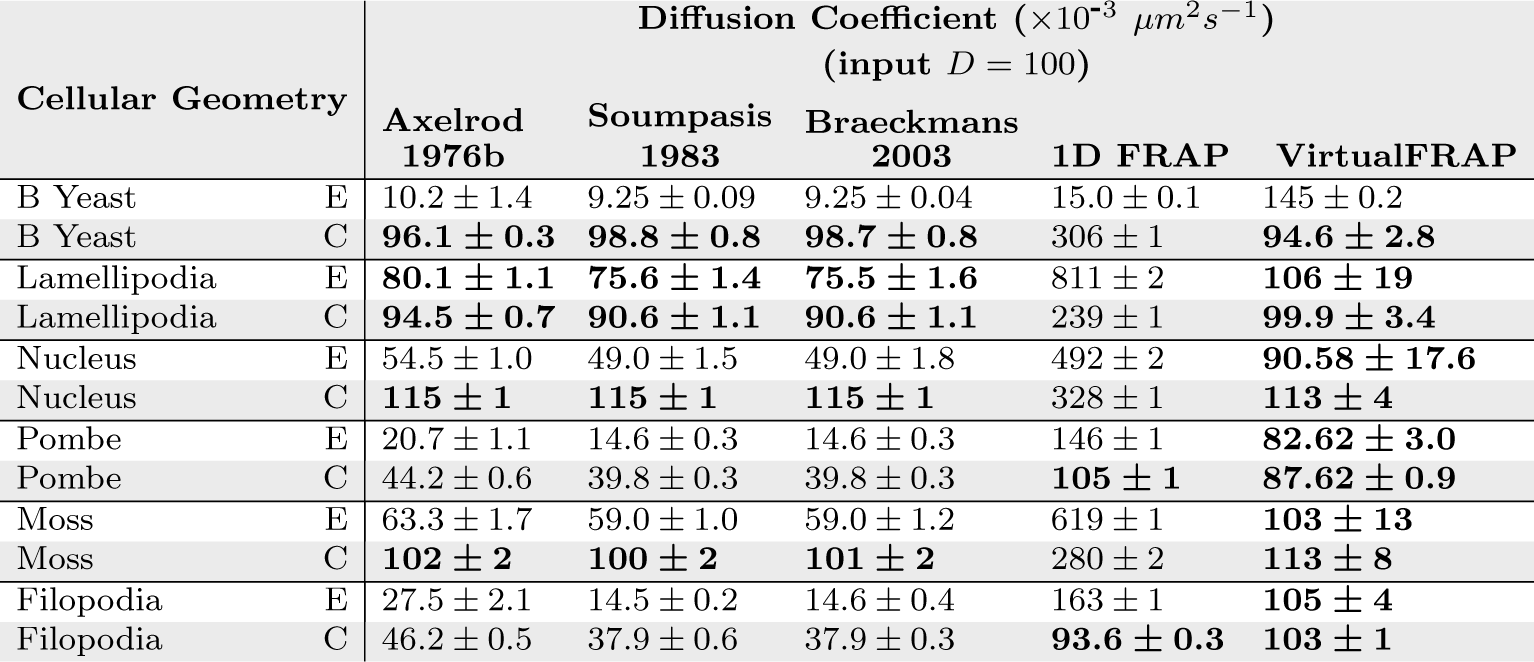
Best fit diffusion coefficients for the four analytical models and the VCell VirtualFRAP tool. Input *D* = 0.1 μm^2^/*s* = 100×10^−3^ *μm*^2^*s*^−1^. ROI is either at the cell Edge (E) or Center (C). Bolded text indicates cases where the model produced an answer within 25% of the true D value. ± represents standard error of ten simulations.

**Table S4.**
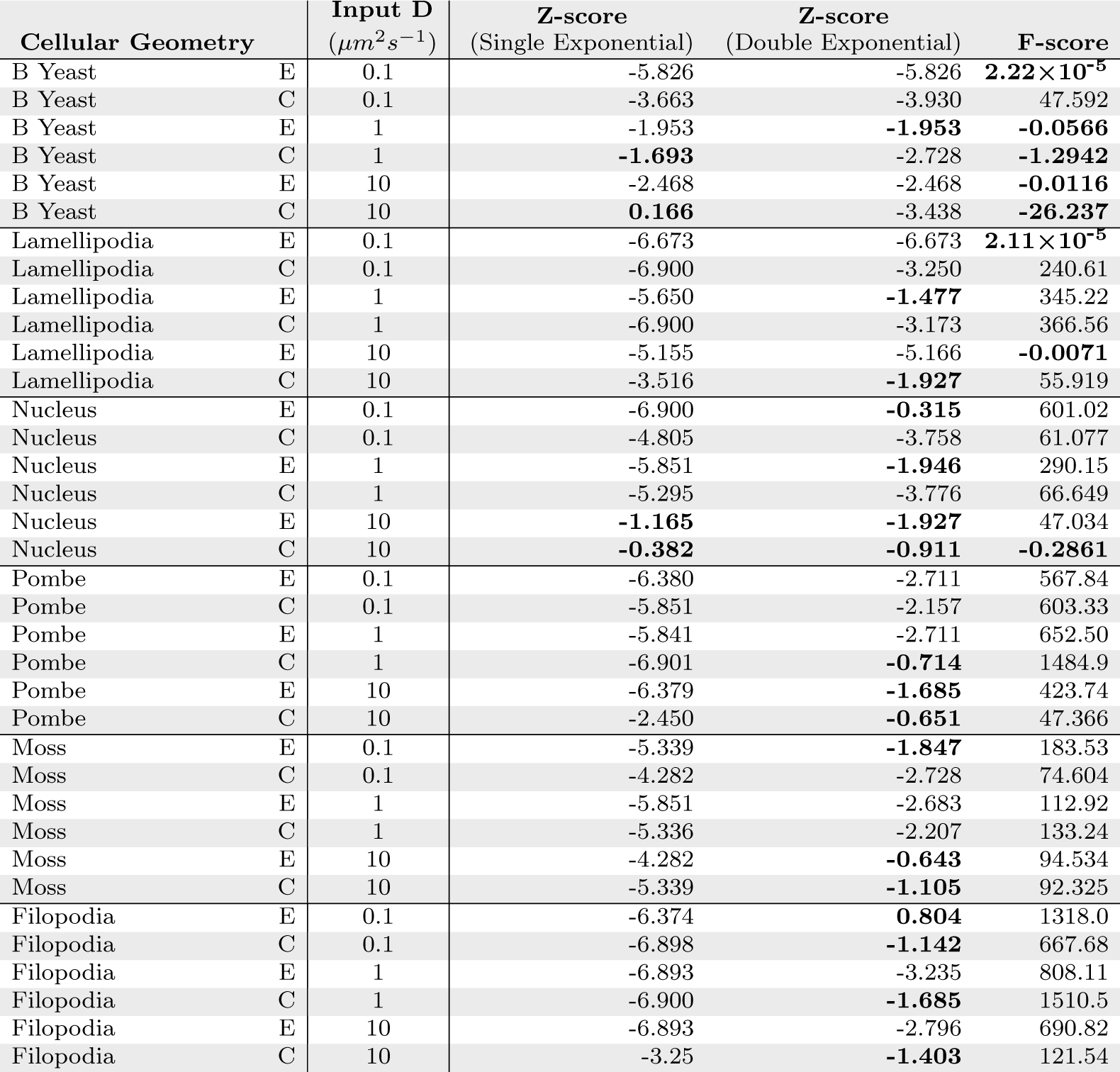
Model selection and evaluation through statistical analysis. Small F-scores indicate that the single-exponential model should be chosen and are denoted with bold text. Large negative Z-scores indicate that the order of the residuals is random (better fit) and are denoted with bold text. ROI location is either at the cell Edge (E) or Center (C).

**Table S5.**
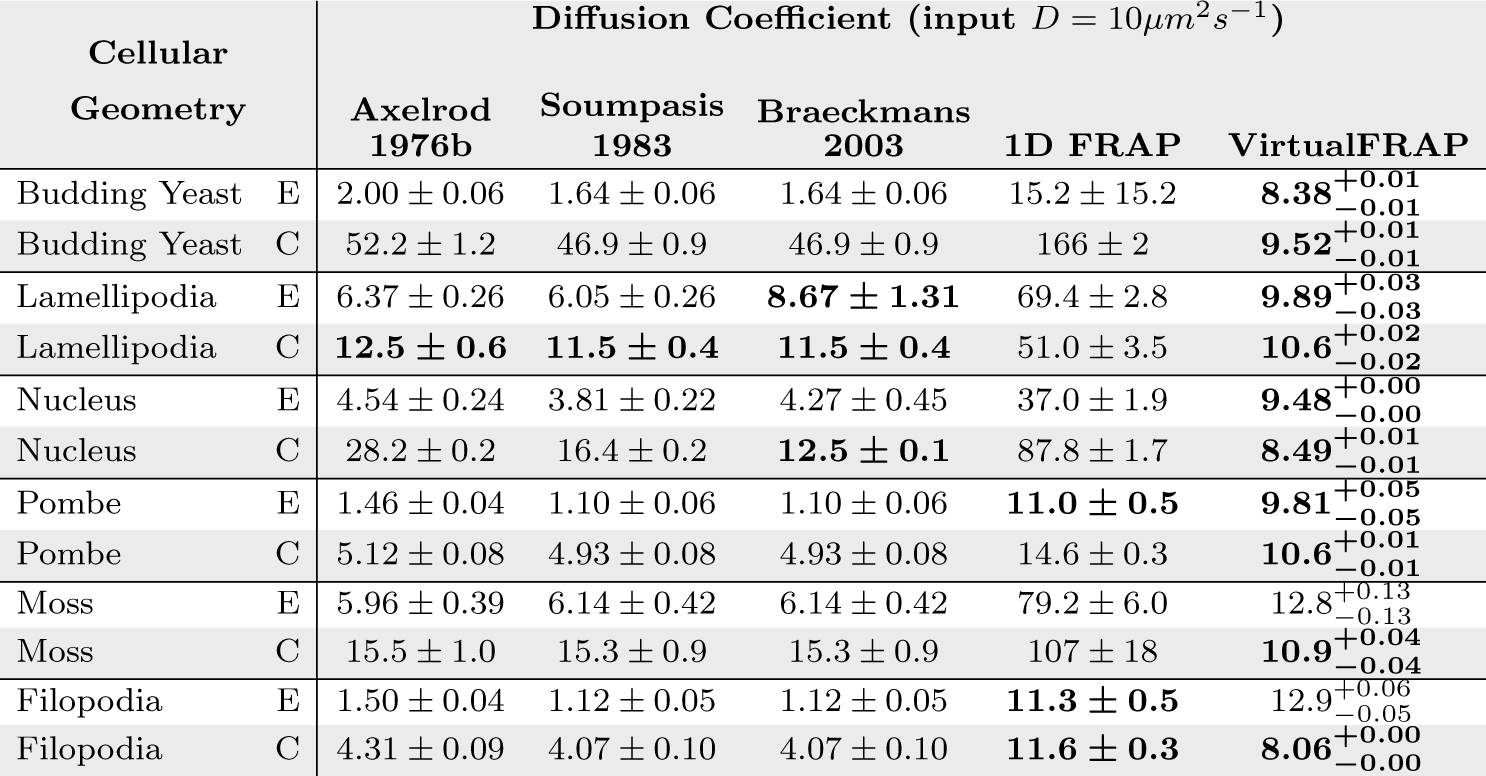
Best fit diffusion coefficients for the four analytical models we used, and the VCell Virtual-FRAP tool. ROI is either at the cell Edge (E) or Center (C). Input *D* = 10 *μm*^2^/*s*, with 280 kHz scan rate. Bolded text indicates cases where the model produced an answer within 25% of the true D value. ± represents standard error of ten simulations for analytical models. For VCell VirtualFRAP, manual use caused us to only use a single replicate; the asymmetric interval returned by the software is presented rather than a standard error.

### S11. Simulated Bleach Profile

The parameters we used for the DCMS simulations produce a bleached region extending all the way through the cells. This is illustrated in **Figure S21** in the moss cell geometry, which is the thickest cell type we tested. This indicates that our bleached region is quasi-cylindrical.

**Figure S21.**
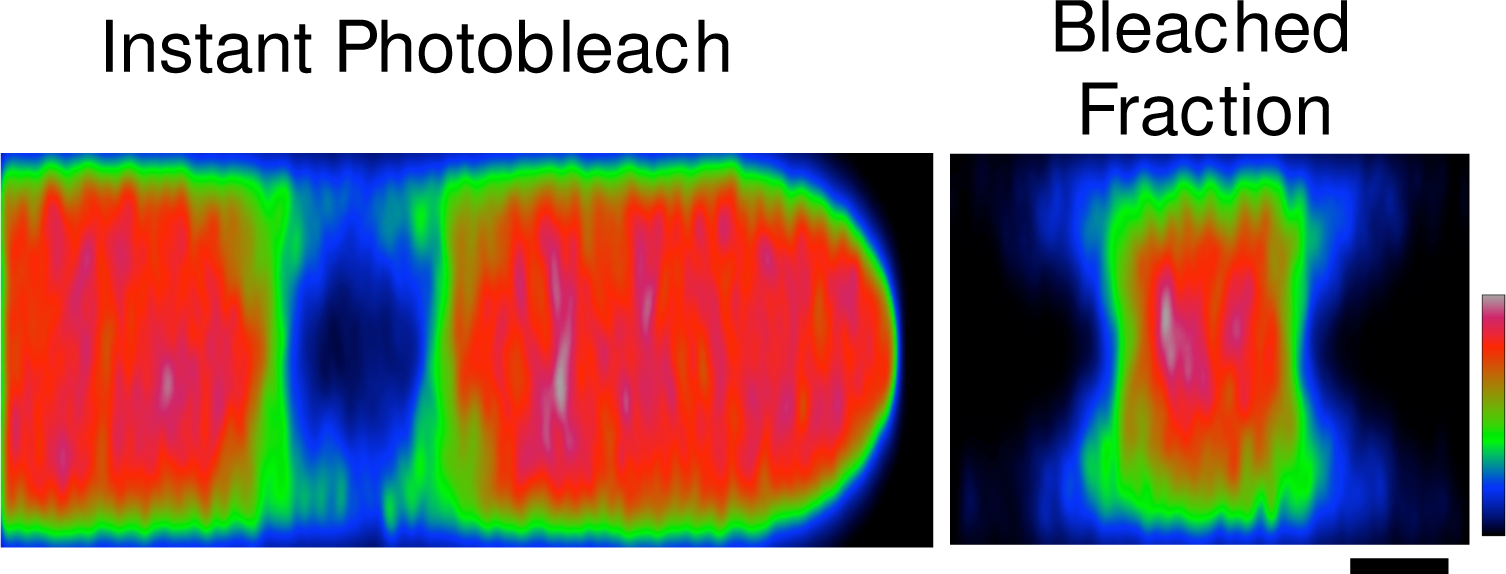
Illustration of the three-dimensional profile of the photobleached region for DCMS moss simulations (left) and the corresponding bleach fraction (right). Immobile fluorophores were used to create a z-stack showing only the effects of photobleaching. Scale bar is 2 *μm*.

### S12. Table of Symbols

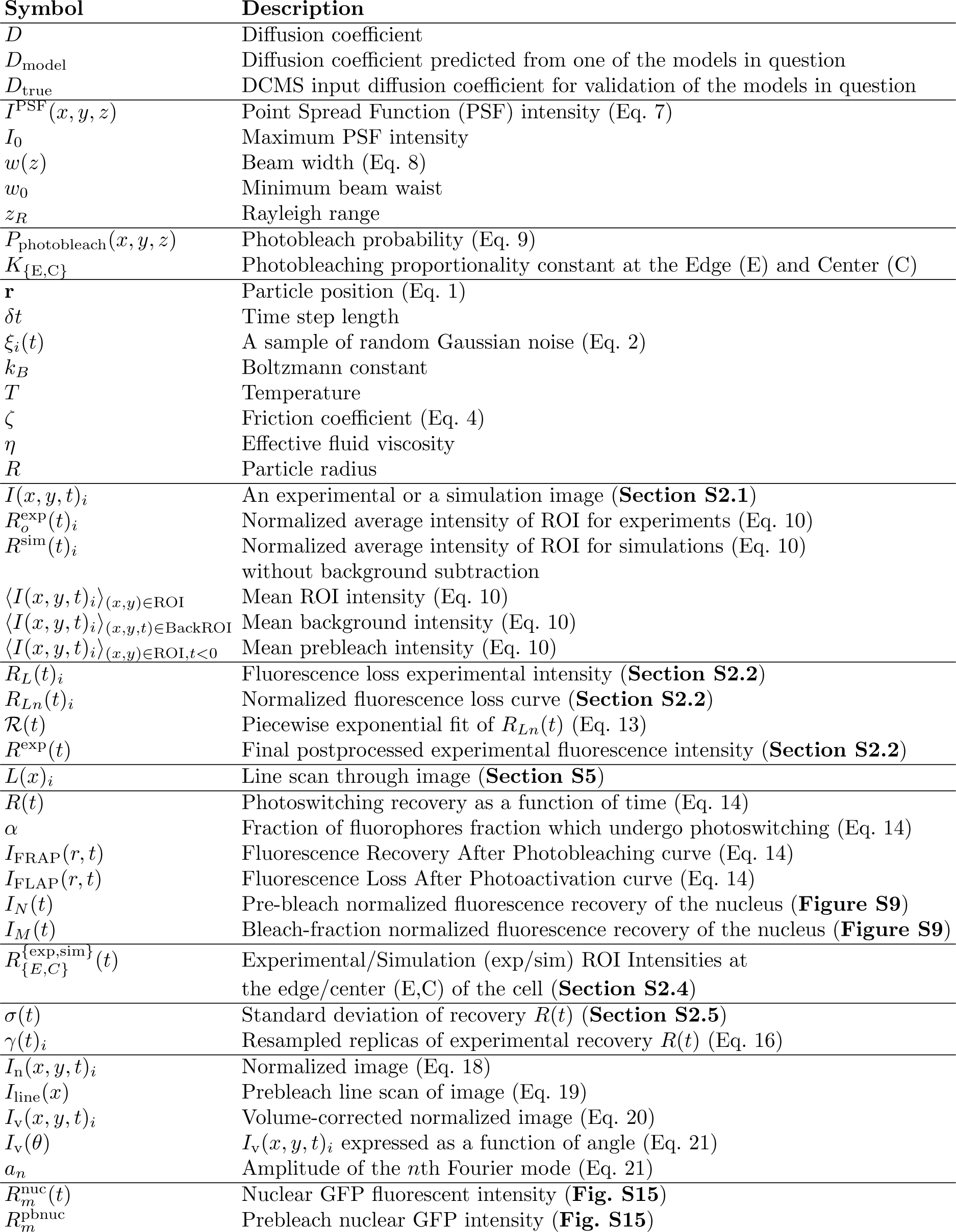

## Supporting Movies

**Movie S1.** Simulation of FRAP at the moss cell edge. Three dimensional (upper left) and two dimensional (upper right) renderings of simulated scanning confocal photobleaching and recovery are shown. Fluorescence intensity is indicated by the rainbow lookup table. Total fluorescence intensity measured in the ROI (4*μm*, located at the edge of the cell) is shown as a function of time, focusing on the overall recovery (bottom) and photobleach (inset). Artificially fast imaging scan rates (instantaneous image capture at 1000 fps) and parameters (*D* = 1.0 *μm*^2^*s*^−1^, 500 × 500 pixel resolution) were used to illustrate three dimensional properties of the simulation.

**Movie S2.** Simulation of FRAP at the moss cell center. Three dimensional (upper left) and two dimensional (upper right) renderings of simulated scanning confocal photobleaching and recovery are shown. Fluorescence intensity is indicated by the rainbow lookup table. Total fluorescence intensity measured in the ROI (4*μm*, located at the center of the cell) is shown as a function of time, focusing on the overall recovery (bottom) and photobleach (inset). Artificially fast imaging scan rates (instantaneous image capture at 1000 fps) and parameters (*D* = 1.0 *μm*^2^*s*^−1^, 500 × 500 pixel resolution) were used to illustrate three dimensional properties of the simulation.

**Movie S3.** Analysis of spatial recovery of 3xmEGFP at the moss cell edge. Cropped and frame averaged photobleaching ROI at the cell edge for 3xmEGFP cell line (left). Corresponding angular intensity profiles of 3xmEGFP at the cell edge (right). The first mode of the Fourier series, *f*_1_, is overlaid in blue, with its magnitude, *a*_1_ shown in the inset. *n* = 14; ROI is 4*μm* in diameter. Mean subtracted image intensity is denoted with the rainbow lookup table.

**Movie S4.** Fluorescence recovery in the budding yeast. Medial confocal slice of simulated fluorescence recovery of budding yeast following photobleaching of the daughter cell is shown. Simulations were run with a diffusion coefficient of *D* = 1 *μm*^2^*s*^−1^.

**Movie S5.** Fluorescence recovery in the nucleus. Medial confocal slice of simulated fluorescence recovery of the nucleus following photobleaching at the cell edge is shown. Simulations were run with a diffusion coefficient of *D* = 1 *μm*^2^*s*^−1^.

**Movie S6.** Fluorescence recovery in the filopodia. Medial confocal slice of simulated fluorescence recovery of filopodia following photobleaching at the cell edge is shown. Simulations were run with a diffusion coefficient of *D* = 1 *μm*^2^*s*^−1^.

**Movie S7.** Fluorescence recovery in pombe. Medial confocal slice of simulated fluorescence recovery of *S.pombe* following photobleaching at the cell edge is shown. Simulations were run with a diffusion coefficient of *D* = 1 *μm*^2^*s*^−1^.

**Movie S8.** Fluorescence recovery in the lamellipodia. Medial confocal slice of simulated fluorescence recovery of lamellipodia following photobleaching at the cell edge is shown. Simulations were run with a diffusion coefficient of *D* = 1 *μm*^2^*s*^−1^.

